# Evolution of ultraconserved elements by indels

**DOI:** 10.1101/2025.05.27.656252

**Authors:** Priscila Biller

**Affiliations:** Physics and Biology Unit, Okinawa Institute of Science and Technology (OIST), 1919-1 Tancha, 904-0495, Okinawa, Japan

**Keywords:** ultraconserved sequences, fragmentation systems, comparative genomics, vertebrate evolution, indels

## Abstract

Ultraconserved elements are segments of DNA that match identically in different species. Finding 100% identity over long evolutionary times is unexpected, but pioneering research in the human-mouse pairwise alignment uncovered something even more puzzling: these elements are not as rare as previously suspected, and their sizes are distributed as a power-law, a feature that cannot be explained by standard models of genome evolution where genome conservation is expected to decay exponentially over time. Despite the power-law behavior having been reported and investigated in a wide variety of biological and physical contexts, from cell-division to protein family evolution, why it appears in the size distribution of ultraconserved elements remains to be fully clarified. To address this question, we propose a model of DNA sequence evolution by mutations of arbitrary length based on a classical integro-differential equation that arises in various applications in biology. The model captures the ultraconserved size distribution observed in pairwise alignments between human and 40 other vertebrates, encompassing more than 400 million years of evolution, from chimpanzee to zebrafish. We also show that the model can be further used to predict other important aspects of genome evolution, such as indel rates and conservation in functional classes.

## Introduction

Evolution of vertebrates is permeated by ultraconserved elements, DNA sequences that have remained unaffected by mutations during millions of years of evolution. Remarkably, most of them are located in non-coding regions, and it is believed that they play a role in important cellular tasks. It has been exactly two decades since the first work describing ultraconserved elements in the human genome was published (Bejerano et al. [2004]), but only more recently have experimental studies been able to thoroughly investigate the reasons for ultraconservation.

Many of these elements were found to play a role in gene expression regulation, in their majority linked to enhancer activity of genes expressed during embryonic development (Pennacchio et al. [2006], Snetkova et al. [2021]). However, the reason for extreme conservation remains a mystery for a significant amount of these elements: In humans, for example, around 17% of long (above 200 base pairs) non-coding ultraconserved elements still have unelucidated function (a recent review can be found in Snetkova et al. [2022]). We hope that the findings reported here will help biologists to target regions of interest for experimental investigation, as experimental functional examination of all regions with unknown functionality is impractical at present.

Due to the limited genomic data available two decades ago, ultraconserved regions were initially defined as exact matches above 200 base pairs in the human, mouse, and rat genomes (Bejerano et al. [2004]). Later, it was confirmed that these sequences also appeared with a high level of conservation in other animals, although they were not always perfectly conserved (Snetkova et al. [2022]). In fact, the number and length of genomic regions identified as “ultraconserved” vary based on the selected species and their divergence time; some studies have broadened the definition to include regions with synonymous mutations or a few mismatches (Cummins et al. [2024]).

Typically, these studies analyze all species at once, with the aim of identifying sequences that are common throughout the entire dataset. In contrast, our goal is to investigate how the number and length of ultraconserved regions change with divergence time, which leads us to focus on perfectly conserved sequences in pairwise alignments of species that diverged at different time points. Due to the phylogenetic connections among species, it is reasonable to expect that evolutionary measurements relying solely on two genomes would be underestimated and less reliable than methods that take phylogenetic dependencies into account. However, as we will elaborate on later, evolutionary distances calculated from only two genomes can achieve a level of accuracy similar to that of estimates based on hundreds of genomes, offering a rapid alternative to comparative analysis that can shorten the estimation time from days or weeks to mere minutes or seconds.

It is also worth noting that most studies on ultraconservation have continued to follow the early approach for identifying ultraconserved elements, ignoring sequences under 200 base pairs. This arbitrary threshold discards the bulk of conserved sequences and, consequently, a tremendous amount of evolutionary signal. By considering both small and large perfectly conserved sequences, we propose a model of sequence evolution to explain the patterns of conservation exhibited in ultraconserved elements. Here we show that these small segments play an important role in the characteristic shape of the size distribution of perfectly conserved regions, and without them, mutation signatures left by the evolutionary process would remain unnoticed.

Interestingly, in terms of their length distribution, the evolution of perfectly conserved elements resembles many other phenomena found in nature, which show a power-law-like behavior in the distribution function of some quantity. To name only a few examples, in physics, the size distribution of meteorites (Hawkins [1964]), and the crushing or grinding of rocks (Domokos et al. [2015]); in chemistry, polymer degradation (Ziff and McGrady [1986]); and, in biology, many genomic properties, such as the sizes of protein families, also have this property (Luscombe et al. [2002]). It is thus of great interest to predict theoretically the evolution of the size distribution during such processes.

A way to mathematically describe these processes is through coagulation-fragmentation equations, a type of integro-differential equations widely used for computing the time-evolution of the amount of particles of a certain size (“particles” can be meteorites, animal groupings or, in our case, conserved sequences). Given their importance in different fields, these equations have been extensively studied and applied in many cases, and explicit solutions have been provided for few classes of model (Ziff and McGrady [1985]) (an in-depth survey can be found in Banasiak et al. [2019]). Only more recently have these equations been used in the context of DNA sequence evolution to try to explain the mechanisms underlying perfectly conserved elements (Massip et al. [2015], Massip and Arndt [2013]). In the current study, we validate for the first time their ability to recover the length distribution of perfectly conserved elements in biological data sets.

We build upon previous research by introducing a unified framework that encompasses various models of sequence evolution within a single formalism. A single parameter can adjust the probability of mutations according to their size, ranging from models that focus solely on substitutions to those where large mutations are more common. This feature enables us to compute estimates for insertion and deletion rates, which are frequently overlooked despite their significance as an evolutionary mechanism for fostering diversity. In humans, for example, the average number of base pairs affected by large mutations per birth is substantially higher than point mutations (see Figure 1 in Campbell and Eichler [2013]; also Kloosterman et al. [2015]).

**Fig. 1:**
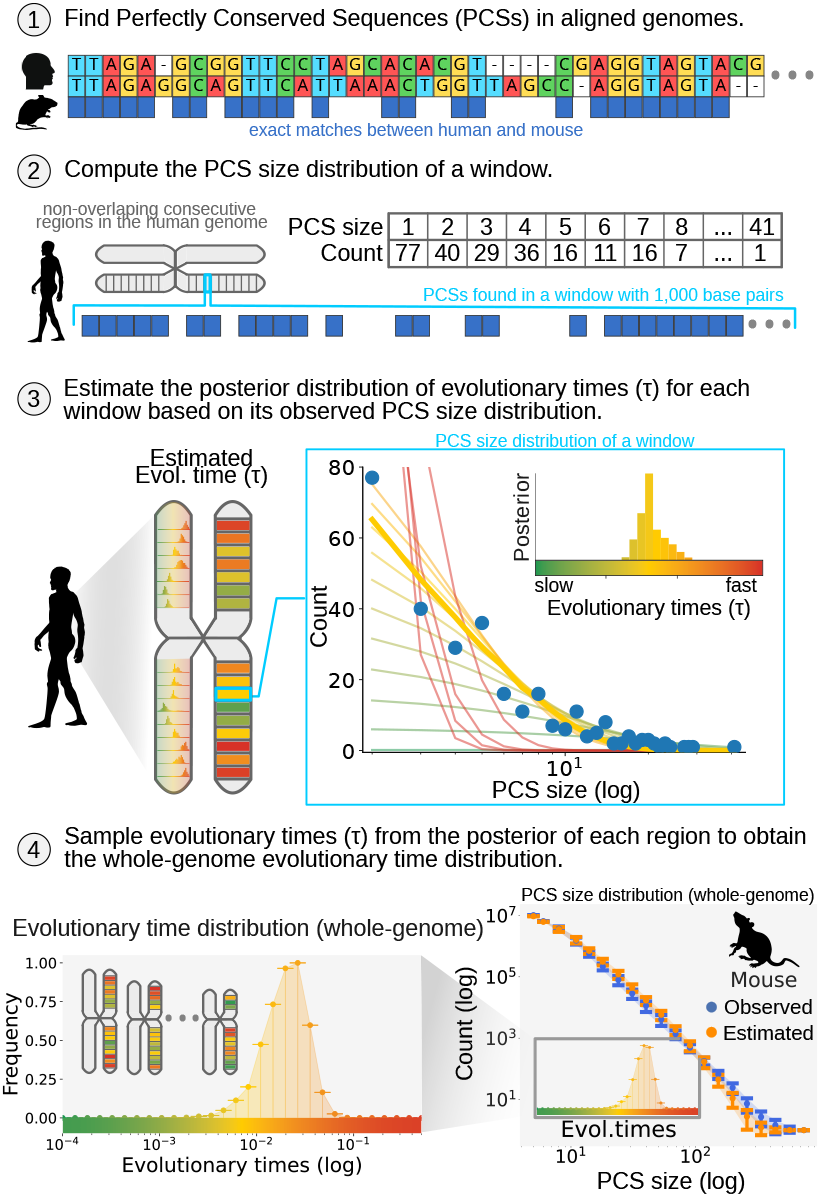
Overview of our method. (1) The human genome is aligned with the genome of another vertebrate. (2) Exact matches (PCSs) between these sequences are extracted and grouped by genomic regions in the human genome. (3) A fitting method is used to compute the posterior distribution of evolutionary times that most accurately explains the PCS size distribution observed in a genomic region. (4) By sampling the posterior distribution of evolutionary times across genomic regions and combining these samples, a whole-genome evolutionary time distribution is defined, which is then used to infer the overall PCS size distribution between the two species.

Another important aspect of the new framework is its ability to estimate evolutionary heterogeneity across different genomic regions. We argue that the power-law-like behavior observed in the length distribution of perfectly conserved elements can be effectively explained by models that consider differences in mutation rates across genomic regions. Despite its simplicity, our model satisfactorily explains the diverse conservation patterns seen in vertebrate evolution and sheds light on which genomic regions have experienced higher or lower evolutionary rates over time.

## Results

### Modeling the evolution of perfectly conserved elements

The evolution of conserved elements can be regarded as a fragmentation process in which a genomic region that is perfectly aligned between two species (no gaps or mismatches) splits into smaller conserved segments every time a mutation occurs. The general kinetic equation for discrete fragmentation is given by Ziff [1992]

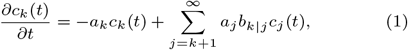

where *c*_*k*_(*t*) is the number of *k*-size particles, or *k*-mers, at time *t, a*_*k*_ is the net rate of breakup of a *k*-mer, and *b*_*k*|*j*_ gives the average number of *k*-mers produced upon the breakup of a *j*-mer.

The first term on the right-hand side represents the decrease in *k*-mers due to all possible breakups of a *k*-mer, while the second term accounts for the gain of *k*-mers by the breakup of particles larger than *k*. The particle size distribution that results from this equation is determined by the details of the kernels *a*_*k*_ and *b*_*k*|*j*_.

Applied to our case, *a*_*k*_ and *b*_*k*|*j*_ encompass all mutations that can occur in a perfectly conserved sequence with *k* base pairs. There are *k* + *s −* 1 positions where a mutation of size *s* can take place, as mutations may be fully contained within a PCS or partially overlap with it. Based on previous findings suggesting an inverse relationship between mutation size and frequency (Levy Karin et al. [2015], Fan et al. [2007], Gu and Li [1995]), we assume that the probability of a mutation with length *s* is proportional to *s*^−*α*^, where *α >* 1 is the slope parameter of the power-law distribution of mutation sizes. If *α* is set to a high value, larger mutations are very unlikely to happen and will play a minor role in evolution; conversely, *α* ≈ 1 gives a prominent role to larger mutations. By taking these observations into account, the above equation can be applied to our case by replacing *a*_*k*_ and *a*_*j*_*b*_*k*|*j*_ by 2((*k −* 1)*ζ*(*α*) + *ζ*(*α −* 1)) and 4*ζ*(*α*), respectively (*ζ*(*α*) is the Riemann zeta function). This integro-differential equation is solvable using the Laplace transform method, yielding the following analytical expression:

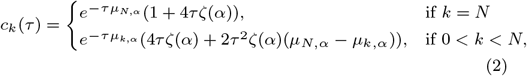

where *N* is the size of the genomic region analyzed (i.e. the “window” size), and *τ* is the evolutionary time; both will be elaborated on in the next paragraphs. A complete derivation of Equation 2 is provided in Appendix A. A model based on the same idea was proposed in an interesting paper by Massip and Arndt [2013], but taking only substitutions into account (*a*_*k*_ = 2*k* and *a*_*j*_*b*_*k*|*j*_ = 4). Under our formalism, their model would be analogous to the special case of choosing a very high value for *α* (Appendix A shows how their exact formula can be obtained by setting *α* → ∞).

A simple method based on Equation 2 can be implemented to compute evolutionary times in different genomic regions. First, the two species of interest are aligned, and their exact matches (PCSs) are identified and grouped into non-overlapping windows. Then, Equation 2 is applied to the PCSs found in a genomic region to evaluate the likelihood of the observed data at a given evolutionary time. By analyzing different evolutionary times, a posterior distribution of evolutionary times can be estimated for each genomic region. Fig. 1 shows a schematic overview of the method, and detailed descriptions of each step are included in the Material and Methods section. Let us briefly discuss the principal assumptions and potential shortcomings of our model.

- *Chronological time vs Evolutionary time*. It is important to emphasize that here time is not treated as chronological time, but as evolutionary time, a dimensionless measure defined as the product of the divergence time (e.g. years) and the mutation rate (e.g. PPPY, per position per year). We are going to use this relation later on to derive mutation rates that can be compared with experimental observations.
- *Irreversible loss of synteny*. By using pure fragmentation equations (no coagulation involved), we are implicitly assuming that once synteny is lost in a genomic region, it cannot be regained. Moreover, for mathematical convenience, we are also assuming that the effect caused by two or more mutations hitting the same spot can be neglected (infinite sites assumption). Depending on the evolutionary time separating two species, these conditions might be more or less likely to hold.
- *Intragenomic mutation rate variability*. To account for rate heterogeneity, the genome needs to be partitioned into consecutive, non-overlapping regions, each of them being assumed to evolve at its own pace. Therefore, the model needs to be applied repeatedly and independently to every region. Although many genomic analysis have used a similar approach— also called “binning” or “windowing”—at diverse scales (e.g. 100kb in Foley et al. [2023] and Chen et al. [2009a]; 25kb in Cooper et al. [2005]; 1kb in our study), caution should be exercised in the choice of the window size. These regions need to be big enough to provide relevant information for the inference, but at the same time as small as possible to avoid mixing fast-evolving and slow-evolving regions. Ideally, the uniform rate assumption should hold inside the region. Concerns regarding the approach of genome partitioning were raised in previous works (Miklós et al. [2004], Siepel et al. [2005]), and will be discussed further below.

### Length distribution of perfectly conserved elements

To validate the model predictions, we used 40 pairwise alignments between humans (hg38) and other vertebrates downloaded from the UCSC Genome Browser. A complete list of their genome assembly IDs and other relevant information can be found in Supplementary Table S1. Perfectly conserved elements were then extracted from pairwise alignments, and their length distribution was compared to those obtained with our model. The estimated PCS size distribution was obtained by sampling the posterior distribution of evolutionary times for each genomic region, and taking these sampled times to derive sampled PCS sizes, with the observed number of PCSs as reference. To assess the variability in the estimates, this process was repeated 100 times and the average estimated PCS size distribution was compared to the observed PCS size distribution, as shown in Fig. 2. Our predictions are in good agreement with biological datasets both at the whole genome level (Fig. 2), as well as at the chromosomal level (Supplementary Fig. S1).

**Fig. 2:**
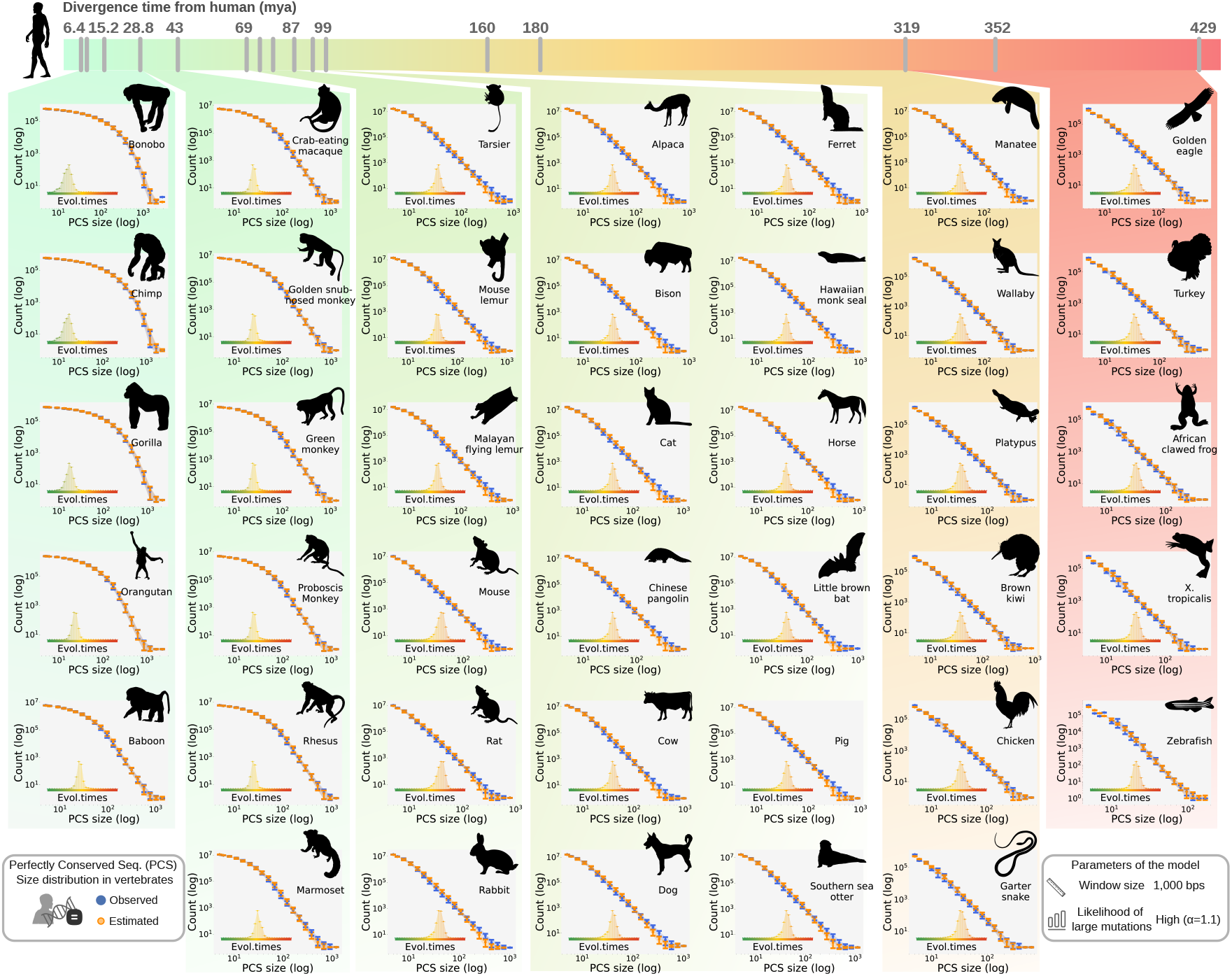
Length distribution of conserved elements in vertebrates. 40 pairwise alignments between human and other vertebrates were used to compute the perfectly conserved sequence (PCS) size distribution across a wide divergence time range. Observed data are shown in blue, and predictions from our model are shown in orange. In the main plots (PCS size distribution), the upper limits for both the x-axis and y-axis vary depending on the maximum PCS size and count found in each species, respectively. To enhance clarity and prevent points from obscuring one another, the x-axis values in the main plot were divided into 20 logarithmic bins, with the y-axis showing the mean PCS count and its standard deviation for each bin. Inset shows the distribution of evolutionary times. The insets of all graphs have the same x-axis (log) and y-axis, both ranging between 0 and 1.

As discussed earlier, our framework can accommodate both large insertions and deletions (*α* → 1), as well as exclusively substitutions (*α* → ∞). Previous works on mammalian datasets (Levy Karin et al. [2015], Fan et al. [2007]) have suggested specific values of *α* for each species, overall varying between 1 and 2, usually closer to 1 than to 2 (Supplementary Fig. S2). Based on these previous findings, we investigated whether large mutations significantly affect the length distribution of ultraconserved elements, by comparing the results of *α* = 1.1 (in accordance with previous works) to *α* → ∞. Remarkably, we find that in both cases, spanning a divergence time range from 10 mya to 400 mya, the length distribution of ultraconserved elements can be predicted with a high level of accuracy and they did not show a significant difference. These findings suggest that variations in *α* can be compensated for by a different distribution of evolutionary times, allowing observed patterns to emerge from different evolutionary processes (Louca and Pennell [2020]). The methods are compared in Supplementary Fig. S3, which complements Fig. 2 by detailing the errors in the estimated versus observed PCS size distributions for each species.

A second important parameter is the window size, which should be kept as low as possible to reduce the variability of evolutionary sequence conservation per region. We therefore chose a window size of around 1,000 base pairs, roughly corresponding to the largest size of a perfectly conserved region observed in the vertebrate dataset (Supplementary Fig. S4). Although previous studies have used considerably larger window sizes, up to 2 orders of magnitude higher than those used here (Foley et al. [2023], Chen et al. [2009a]), we noticed that, at least in our case, a large window size can hide information from slow-evolving areas, leading to an overestimated evolutionary time. This can be explained by the relation between number of conserved elements and evolutionary time: as time increases, conserved elements become smaller and more numerous; thus if a region contains both high and low evolutionary times, the overall conserved length distribution will be dominated by segments with higher evolutionary time (i.e. the fast-evolving regions). This point is addressed in more details in Appendix C.

In this work, we present for the first time, to our knowledge, a comprehensive analysis of the time-evolution of lengths of conserved elements over a wide range of time scales. We observe that these distributions first take on an exponential-like shape (Fig. 2, top left corner), and then shift to a heavy-tail form after a sufficiently long evolutionary process (around 30–40 mya). Notably, Luscombe et al. [2002] found the exact same behavior in the distribution of gene families sizes. When trying to determine whether a power-law or another function would be the best fit for this type of data, they observed that a triple-exponential showed the smallest residual among all candidate functions, and probably higher-order exponentials would provide increasingly better fits. However, they could not provide a reason for choosing this function over the other options (lognormal and power-law distributions also performed well). In the context of ultraconserved elements, our findings provide a compelling rationale for the use of the best-fitting option: If evolution is regarded locally, an exponential-like distribution should be observed; the heavy-tail behavior arises from the variation of these exponents throughout the genome, and can be regarded as a sign of rate heterogeneity. Thus, our model reconciles apparent inconsistencies between power-law empirical observations and exponential-like predictions from theoretical proposals.

### Evolutionary time: pairwise-based estimates vs. phylogenetic approaches

In addition to the length distribution of conserved sequences, it would be ideal if the estimated evolutionary times could also be used to provide accurate predictions about other aspects of the evolutionary process. However, since our estimates are based on pairwise comparisons, we must first assess whether this could result in an underestimation of evolutionary times due to a lack of phylogenetic constraints. If the differences between two species do not provide sufficient information to infer their evolutionary distance, how many additional species would be needed for an accurate estimate?

To investigate this question, we compared our pairwise estimates with phylogeny-based estimates obtained from large phylogenetic trees recently published (Kuderna et al. [2023], Upham et al. [2019]). The first tree analyzed comes from Kuderna et al. [2023] and encompasses 233 primate species, of which 15 are present in our dataset of 40 vertebrate species. Their tree was inferred with IQ-TREE2 (Minh et al. [2020]), and branch lengths were estimated with the GTR+F+R7 model, a substitution-only model that accommodates different rates of change for each nucleotide pair, and uses a discrete gamma distribution to account for variation in substitution rates across sites. The second tree, from Upham et al. [2019], depicts the evolutionary history of over 5,000 mammals. Branch lenght information was obtained with RAxML (Stamatakis [2014]) and the GTRCAT model. Our dataset includes 31 species that overlap with their dataset.

Fig. 3 shows a comparison of evolutionary estimates between our method and phylogenetic-based approaches. For a given species shared between our dataset and theirs, we computed the mean and standard deviation of the estimated evolutionary time distributions (shown in the insets of Fig. 2) and compared these values to the sum of the branch lengths along the path connecting humans to the given species in their phylogenetic tree. In an ideal scenario, we expect that these values should be roughly the same, as both represent the product of chronological time and substitution rate. For a correct comparison, we have to consider the DNA regions analyzed in each study. Kuderna et al. [2023] focused solely on genomic regions with “large” PCSs (above 50 bps), while Upham et al. [2019] included the entire DNA sequence in their analysis. Therefore, compared to the windows analyzed in our work, the first dataset is a subset that includes only windows with large PCSs, while the second dataset is a superset that contains windows that were excluded for lacking PCSs (Fig. 3, panel A).

**Fig. 3:**
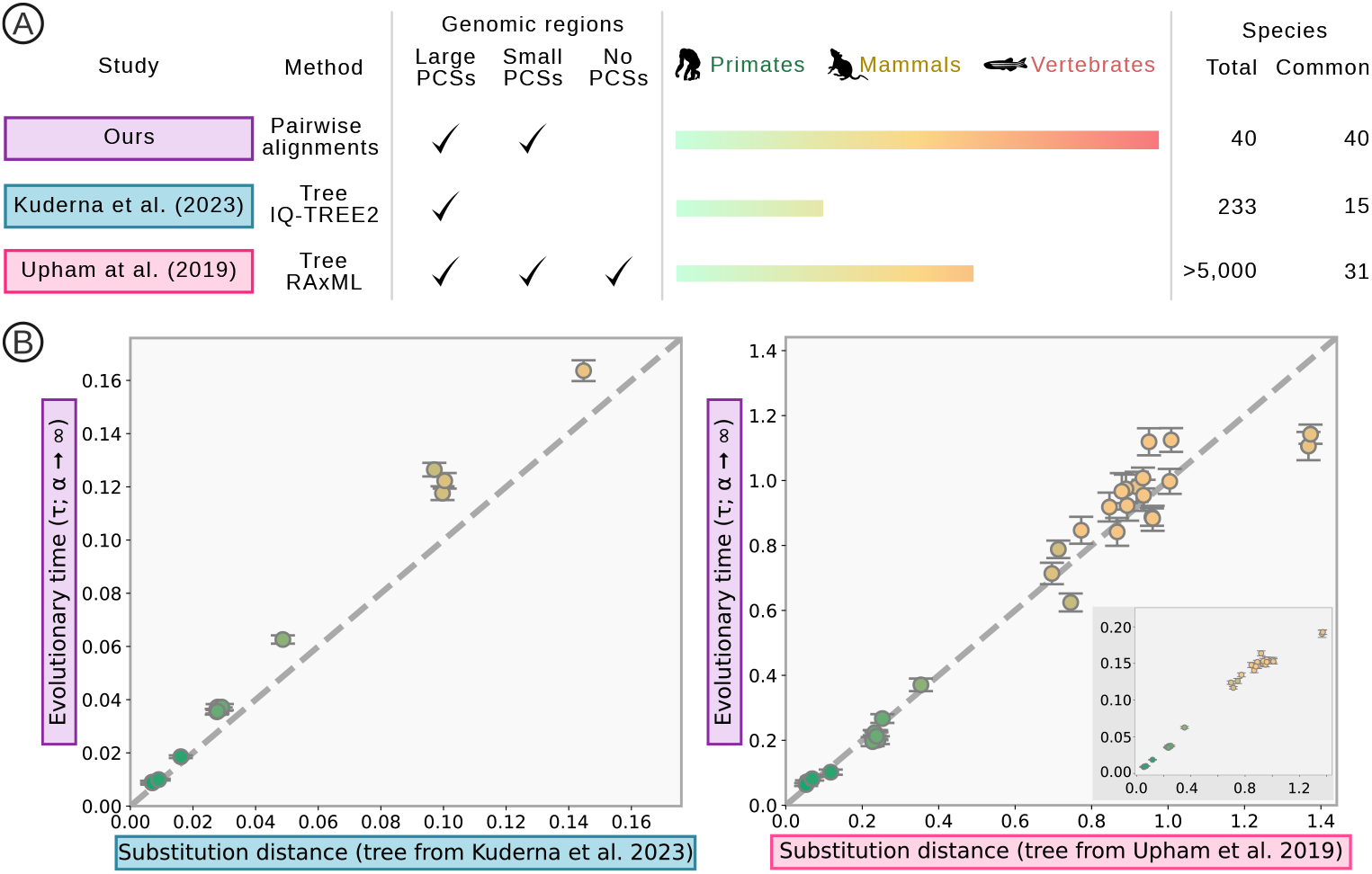
Comparison of evolutionary estimates based on pairwise alignments (ours) versus those based on phylogenetic analysis (Kuderna et al. [2023] and Upham et al. [2019]). *Panel A* shows an overall comparison of the datasets used in the study. Column *Method* specifies whether the evolutionary distances are based on phylogenetic trees or pairwise comparisons. Column *Genomic regions* specifies which DNA segments were included in the analysis: regions with PCSs exceeding 50 base pairs are labeled as “Large PCSs,” those with PCSs above 5 base pairs but below 50 are termed “Small PCSs,” and regions lacking PCSs or with PCSs under 5 base pairs are classified as “No PCSs.” Column *Species* outlines the types of species included in each analysis, with *Total* representing the overall number of species and *Common* denoting the number of species shared with our dataset. *Panel B* presents a comparison of our estimates (y-axis) with those from other studies (x-axis). Each dot corresponds to a species shared by both datasets, with the color of the dot indicating its divergence time from humans, following the color code from panel A (green for primates, yellow for mammals, and red for vertebrates). In the right graph, our estimates have been modified to account for windows lacking PCSs, with the original values shown in the inset plot.

Due to these differences in the dataset, our evolutionary estimates for the mammalian dataset, although well correlated with Upham’s, do not have the same absolute values (Fig. 3, panel B, inset of the right graph). If we also take into account the number of windows without PCSs in each species and assign a fixed high mutation rate to them (see Appendix D), the estimates also match in absolute values, as shown in Fig. 3, panel B, right graph. Regarding the evolutionary estimates from the primate dataset of Kuderna et al. [2023] (Fig. 3, panel B, left graph), we found that our estimates are nearly identical to theirs in absolute values, albeit slightly overestimated, which is expected as they only consider windows with large PCSs.

Even though our approach relies on only two genomes rather than a larger dataset with thousands of genomes, our method’s evolutionary distances show a good match with those reported in the other two studies. Considering the often prohibitive computational costs of analyzing thousands of genomes at once relative to the accuracy improvements it may offer, our method provides a significantly faster and yet reasonably accurate alternative to them.

### Indel rates across vertebrates

Most phylogenetic analyses today are carried out with evolutionary models centered exclusively on substitutions. Phylogenetic trees found in the literature, including those of Kuderna et al. [2023] and Upham et al. [2019], are based on popular bioinformatics tools like RAxML (Stamatakis [2014]) and IQ-TREE2 (Minh et al. [2020]). The core of these tools is the GTR model, an evolutionary model in which substitution rates vary depending on the pair of nucleotides considered.

Larger mutations, such as insertions and deletions (indels), are not considered by these models, despite being one of the most common sources of genetic variation (Mills et al. [2011], Chen et al. [2009b]). Indels are a class of mutations separate from base substitutions, differing in how they originate. The molecular mechanisms underlying indels are reviewed by Savino et al. [2022], including, for example, misaligned sequences during DNA replication (strand slippage) or problematic repair of double-strand breaks.

Probabilistic-based models for indels are far less developed compared to substitution models, with limited options available (Miklós et al. [2004], Lunter [2007]) and recent attempts to make them more efficient (Loewenthal et al. [2021], Levy Karin et al. [2017]). By using an alternative approach centered on the lengths of conserved sequences, our framework can provide evolutionary estimates based on insertions and deletions with a fast analytical equation that simplifies computation.

In this section, we compare our model predictions for indel rates with those from previous studies. Due to the lack of phylogenetic studies based on indel models, and the computational cost of running indel-based models on a large dataset, we need to resort to other types of analyses from earlier studies. A way to assess indel rates is using the relation *µ* = *τ/*2*t*, where *µ* is the average indel rate on the lineage connecting two present-day species, *t* is the divergence time between them, and *τ* is the evolutionary time derived from observable differences in their genomes. This method of computing mutation rates is called “indirect estimation” (Scally and Durbin [2012]), and was used in previous studies based on pairwise alignments (Lunter [2007], Nachman and Crowell [2000]). In our case, each pair of species has a distribution of evolutionary times, so we use the mean value of this distribution as *τ*, whereas the divergence time *t* comes from previous studies (mostly TimeTree Kumar et al. [2022]; see complete information in Supplementary Table S1.

A more direct approach consists of using families to look for mutations carried by a child but not by either of their parents. This type of analysis is conducted more commonly in human populations (Maretty et al. [2017], Besenbacher et al. [2016], Kloosterman et al. [2015], Palamara et al. [2015], Besenbacher et al. [2015], Kondrashov [2003]); however, since our dataset focuses on comparisons between humans and other vertebrates, direct measurements of the indel rate in modern humans provide a valuable reference point for our study.

Fig. 4 shows the comparison of our estimates with the indel rates obtained from both direct and indirect methods (Maretty et al. [2017], Besenbacher et al. [2016], Kloosterman et al. [2015], Palamara et al. [2015], Besenbacher et al. [2015], Kondrashov [2003], Lunter [2007], Nachman and Crowell [2000]); some adjustments and corrections were necessary to make them comparable (more details in Supplementary Table S2). It is important to note that the maximum mutation rate that can be measured with our method decreases as divergence time increases. For instance, with the indel model used for our estimates (*α* = 1.1), we expect that genomic regions with an evolutionary time above 0.10 will not provide enough information for inference, due to an insufficient number of PCSs (Supplementary Fig. S6, panel A). For the human-chimp lineage, this implies that if mutation rates exceed 413.22 *×* 10^−11^ PPPY (per position per year), our method can only offer a lower bound; meanwhile, for the extreme human-zebrafish lineage, the maximum measurable mutation rate is limited to 11.65 *×* 10^−11^ PPPY (see Supplementary Fig. S6, panel B for other species). This constraint does not pose a problem when we focus exclusively on conserved regions, as their mutation rates remain low. However, to ensure that our findings are comparable to those of previous studies that considered the entire DNA, we also had to consider regions without PCSs. Thus, we expect that for species that have diverged over a considerable time (shown in the red zone of Fig. 4), our method provides a conservative estimate of the actual mutation rate.

**Fig. 4:**
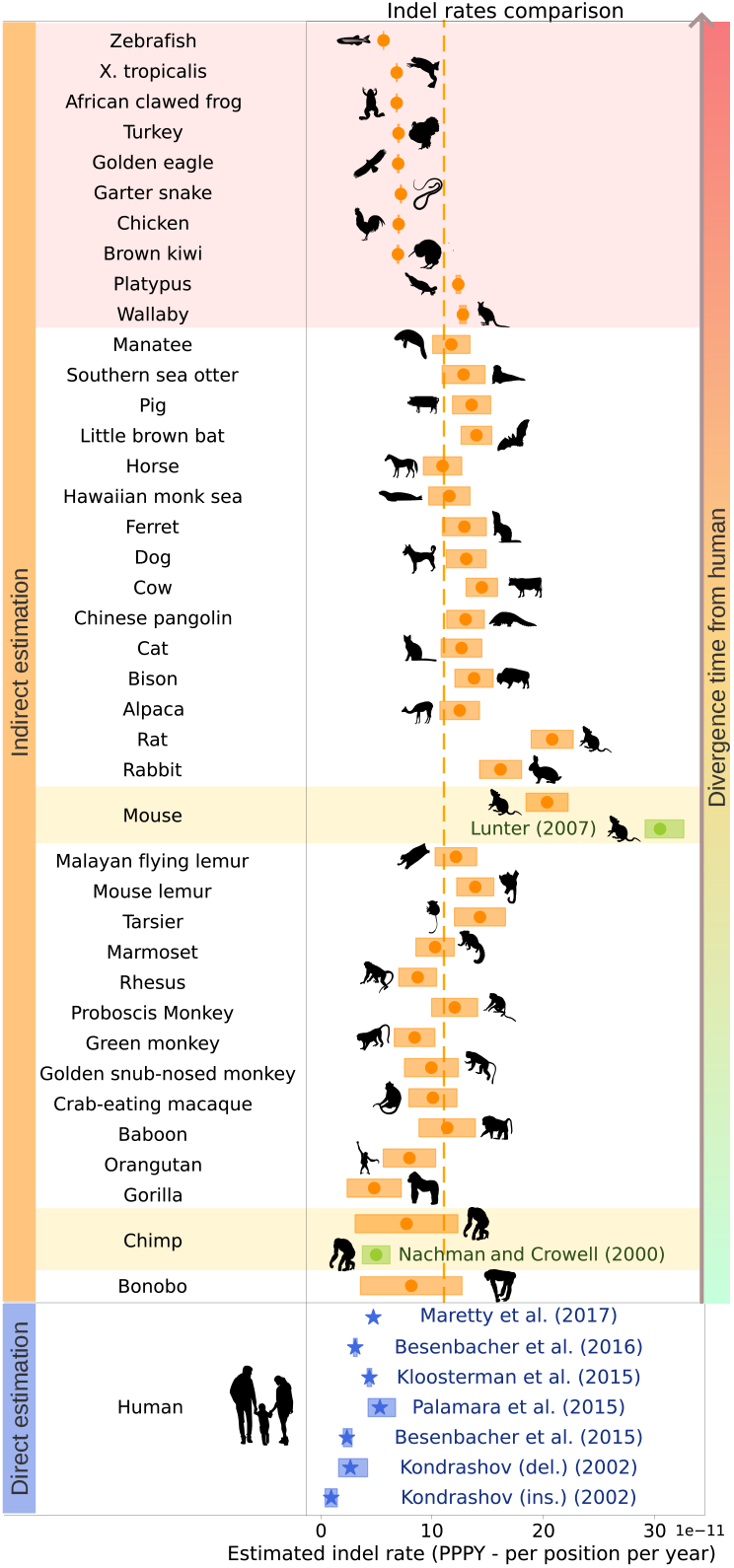
Indel rates in the lineage relationships between human and 40 other vertebrates. “Direct estimates” (blue) refers to methods that count mutations that occur between generations in present-day individuals, whereas “Indirect estimates” (orange and green) refers to estimates based on the evolutionary distance separating two species divided by (twice) their divergence time. Our estimates are shown in orange, with the rectangle borders representing the standard deviation, and the middle point showing the mean indel rate. Indirect and direct estimates from previous studies are indicated in green and blue, respectively. All indel rates were adjusted to “per position per year” (PPPY) in order to make them comparable. The orange dashed line indicates the average indel rate across all species. If evolution were uniform across all lineages, values should be concentrated around this point.

An interesting observation is that, if evolution were occurring uniformly across all lineages, we would expect all indel rates to cluster around the mean value, shown as an orange dashed line in Fig. 4. Nevertheless, as extensively documented in the literature (Hodgkinson and Eyre-Walker [2011], Lynch et al. [2016]), mutation rates can vary significantly depending on the clade in question. For example, the evolutionary paths between humans and other Homininae (Gorilla-Homo-Pan) have indel rates comparable to the ones observed in modern human populations (blue estimates in Fig. 4); however, compared to other vertebrates, these evolutionary rates appear to have slowed down. Conversely, indel rates are particularly high in the evolutionary paths of humans and rodents (rats and mice). Lunter [2007] reported an indel rate of approximately 30.46 *×* 10^−11^ PPPY, while our estimate stands at 20.04 *×* 10^−11^ PPPY, both of which are significantly higher than the average indel mutation rate observed in vertebrates (as a reference point, although not directly comparable to the average mutation rate along the entire evolutionary path, the estimated indel rate in extant wild mice is about 31 *×* 10^−11^ PPPY (Uchimura et al. [2015])). This variation might be attributed in part to a “generation-time effect” (Li et al. [1996]), in which evolutionary rates appear to accelerate in chronological time for species with a shorter generation time, since they go through more generations per time unit. For example, generation times in humans and mice are estimated to be around 27 years (Wang et al. [2023]) and 9 months (Lindsay et al. [2019]), respectively. Previous studies indicate that mutation rates are more comparable among organisms when measured in terms of generation (per position per generation: PPPG) than in terms of absolute time (per position per year: PPPY), but this approach introduces an additional problem of estimating how generation times vary through vertebrate evolution, which is out of the scope of this article (for an in-depth investigation of this problem, we refer the reader to Bergeron et al. [2023], Evans et al. [2012]).

### Functional classes in slow-evolving regions

We next aimed to visualize and compare the evolution of major functional classes, such as protein-coding regions, gene regulatory regions and repeats. We are particularly interested in quantifying the proportion of these classes overlapping regions identified by our method as slow-evolving, and whether this information varies as divergence time increases.

Identifying functional classes in slow-evolving regions requires two steps. The first step is to create a map between the functional classes and the windows used in the previous analysis based on well-known genome annotation databases (GENCODE-V44, RefSeq, RepeatMasker), which is described in further detail in Materials and Methods. An annotation is mapped to a window if the annotation coordinates and the window coordinates intersect. Thus, a single window can be mapped to several functional classes, and the size of the overlap (in base pairs) is taken into account in our analysis. The second step consists of selecting windows based on whether or not they are slow-evolving. Here we used the estimated evolutionary time distribution (shown as insets in Fig. 2) to select windows according to the position of its estimate within the overall distribution. Notice that the selected windows may differ between pairwise alignments, as a window might appear conserved in one instance but less conserved in another. In the results that follow, we take the slowest-evolving windows whose combined sizes sum to 250 MB. This cutoff value is based on a previous work by Rands et al. [2014] suggesting that around 250 million base pairs (250 MB) in the human genome— approximately 8% of the entire genome—are presently subject to purifying selection. Inspired by this, we sought to identify which human functional elements compose the top 8% most conserved regions in human lineage over time.

Fig. 5 presents the overlap between slow-evolving regions and functional classes, across various functional classes (rows) and throughout vertebrate evolution (x-axis, common to all graphs). The y-axis shows the percentage of slow-evolving regions that are part of a specific functional class, with the baseline being defined as the proportion of the functional class in the whole-genome (solid black line in the center of each plot). Results higher (lower) than the baseline indicate an over (under) representation of the functional class in slowly evolving regions compared to the whole-genome. Each point on the x-axis corresponds to a specific pairwise alignment between human and another vertebrate, sorted by divergence time (from the species closest to human on the left to the farthest species on the right).

**Fig. 5:**
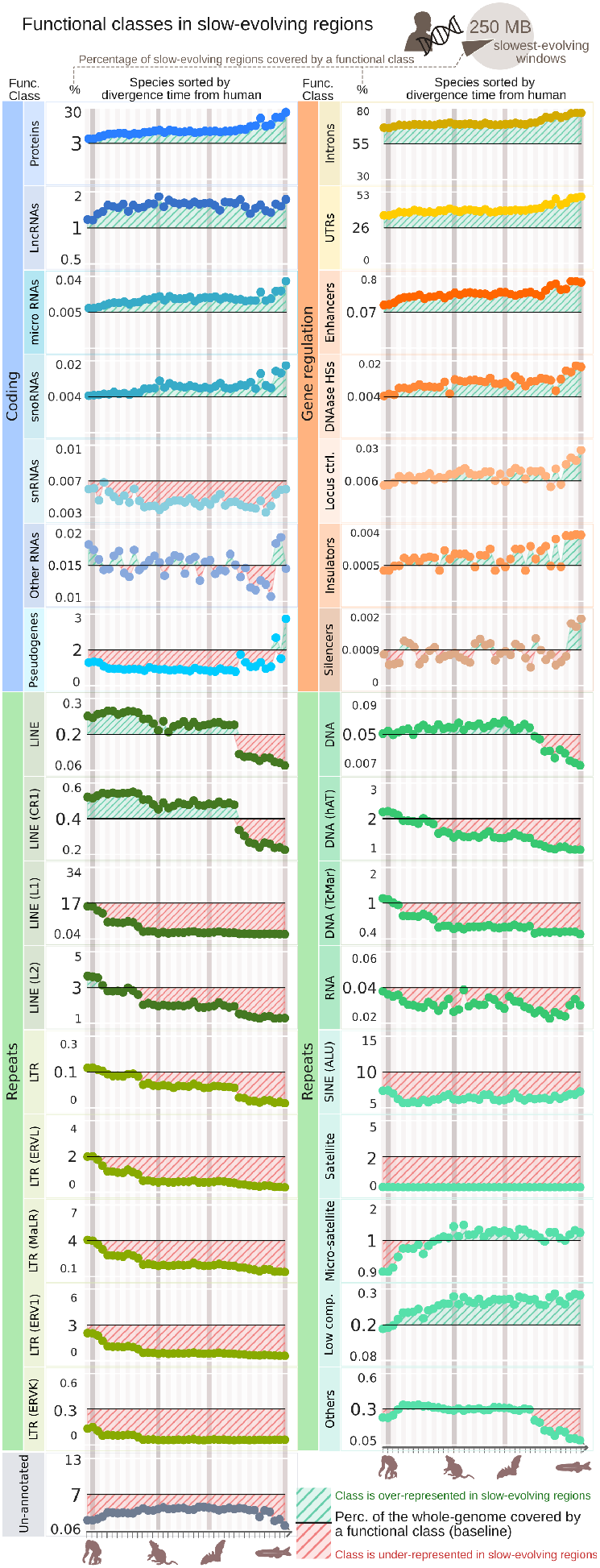
Distribution of slow-evolving regions across functional classes. Rows indicate different functional classes, columns indicate different pairwise alignments between human and another vertebrate (left: closest species to human; right: farthest species from human). For each pairwise alignment, the set of “slow-evolving” windows was defined by selecting the windows with the lowest evolutionary times, summing up to 250 MB (around 8% of the human genome). For each functional class, the y-axis shows the percentage of windows identified as “slow-evolving” that fall under that class, compared to the baseline (solid black line in the middle of each graph), which denotes the percentage of windows in the entire genome under the same functional class.

In total number of base pairs, slow-evolving regions are overwhelmingly dominated by regulatory regions directly adjacent to coding regions. On average, 69% and 40% of slow-evolving regions are annotated as introns and untranslated regions (UTRs), respectively. However, if we consider the contribution from each class compared to how frequently they are found in the whole genome, other classes appear more represented in slow-evolving regions than introns and UTRs.

Confirming previous findings (Pennacchio et al. [2006]), enhancers are the most over-represented elements in slow-evolving regions across the vertebrate evolution (on average, they appear 5.2 times more often in slow-evolving regions than in the whole-genome). Special attention in recent years has been given to these ultraconserved enhancers (Wong et al. [2020], Snetkova et al. [2021]). To date, the literature is inconclusive on whether or not their extreme degree of sequence conservation is maintained by strong purifying selection (Wong et al. [2020], Snetkova et al. [2021], Ahituv et al. [2007]). Despite methodological studies having used low mutation rates as a proxy of purifying selection (Rands et al. [2014]), in vivo studies observed that enhancers can still keep their function even without perfect conservation (Wong et al. [2020], Snetkova et al. [2021], Ahituv et al. [2007]) (although see Dickel et al. [2018] for a counterpoint), and similar sequence conservation can show apparent differences in conservation of activity (Villar et al. [2015]). Apart from enhancers, DNase I hypersensitive sites are another main gene regulatory group that consistently shows a significant presence in slow-evolving regions, appearing, on average, twice as often as in the whole genome.

Notably, few repeat families also appear overrepresented in slow-evolving regions. LINE (Long Interspersed Nuclear Elements) repeats and low-complexity repeats have a significant overlap with slow-evolving regions (1.24 and 1.28 times more frequently than in the whole genome, respectively), similar to introns (1.26 times). Unlike gene regulatory elements, their contribution varies throughout vertebrate evolution. For example, while low-complexity repeats are shared between humans and other species back further in evolution, LINE repeats are mostly shared up to 100 mya. These findings indicate that, despite being commonly grouped together, repeat families display distinct conservation patterns, with conservation within these families often being neglected. A similar observation was made by a previous study on ancient repeats (Kamal et al. [2006]), which found a large family of ancestral repeats under strong selection despite ancestral repeats being frequently used as proxies for neutral evolution (Lunter et al. [2006]).

## Discussion

Growing interest in ultraconserved elements (Dickel et al. [2018], Snetkova et al. [2021, 2022], Cummins et al. [2024]) reinforces the importance of developing a solid theoretical framework to guide experimental research on the topic. Here, we introduce a model that describes the evolution of perfectly conserved sequences in the genome, allowing for adjustments to focus exclusively on substitutions or on insertions and deletions of different lengths, with the potential for future adaptations to other evolutionary mechanisms.

Our findings represent a significant step toward establishing the much-needed theoretical framework for experimental studies. For instance, one poorly understood phenomenon (Salerno et al. [2006]) was the striking regularity found in highly conserved sequences, where their length and frequency are linked by a power-law, a pattern that classical models of genome evolution, which expect an exponential distribution, are unable to explain. Our model enabled us to show that power-law behavior is the result of regions evolving at different rates, combined with enough time to let these differences accumulate and become noticeable. Thus, the apparent paradox can be reconciled if classical models of genome evolution are applied locally instead of globally, where the evolutionary process does not deviate significantly from the assumption of a uniform evolutionary rate.

A notable characteristic of our model is its capability to provide meaningful insights into the evolutionary history of two species using exclusively their genomic data, eliminating the need for additional genomes or other sources of information. In a time where evolutionary analyses are conducted on thousands of genomes (Upham et al. [2019], Kuderna et al. [2023]) and require substantial computational resources, this raised the question of how our estimates compare to those derived from much larger datasets. In the context of inferring evolutionary distances, our estimates closely match those derived from well-stablished substitution models in larger datasets, indicating that the differences observed between two sequences can yield valuable information when properly considered.

Importantly, our model enables us to examine the evolution of insertions and deletions, a process that is not typically implemented by popular bioinformatics tools and often excluded from phylogenetic analysis. We offer a first demonstration of this feature by providing estimates for the indel rates of all lineages linking humans with other 40 vertebrates, along with an analysis of the functional classes found in ultraconserved sequences and the temporal changes in their distribution.

The ability to have a fast analytical solution to the various evolutionary models encompassed by our framework opens up the possibility of investigating many interesting issues related to ultraconserved elements. One of them is to assess the differences in purifying selection acting on indels versus substitutions in various genomic regions. In protein coding regions, for example, if the number of nucleotides added or deleted is not a multiple of three, the reading frame is shifted, likely resulting in fitness loss. Even when indels are in frame, they accumulate unevenly along the protein sequences due to their three-dimensional structure (Guo et al. [2012]). Protein structural elements have varying degrees of tolerance to insertions and deletions, with unstructured regions, such as loops, allowing for indels more frequently (Simm et al. [2007], Guo et al. [2012]). Although previous research has identified several factors that contribute to differences in purifying selection between indels and substitutions, the variations seen in protein-coding regions are not yet fully understood. Research in yeast and fruit flies shows significant variation depending on the specific protein, with the purging rate for indels sometimes reaching up to 100 times higher than that of point mutations (Tóth-Petróczy and Tawfik [2013]), which cannot be fully explained by the need to maintain the reading frame.

The selective pressures acting on indels and substitutions in other genomic regions, such as enhancers or specific families of repeats where ultraconserved regions are concentrated, have been even less explored. The estimates provided by our model could help answer these questions in future work. A potential initial step is to look into the substitution and indel rates that have already been computed for the vertebrate dataset, and assess the variation in the ratio of nucleotide substitution and indel rates across different genomic regions. Other questions for future research include exploring whether the evolutionary times derived from an indel-based model could help to solve conflicting phylogenetic relationships that arise from substitution-based models (other evolutionary processes, such as introgressions and horizontal gene transfers, have proven useful for this purpose (Coleman et al. [2021], Fontaine et al. [2015])), as well as conducting a more thorough analysis of unannotated regions classified as ultraconserved by our method, which could lead to the identification of new functional elements.

## Methods

### Identification of Perfectly Conserved Elements in Vertebrates

Whole genome pairwise alignments were downloaded from the UCSC Genome Informatics website (http://genome.ucsc.edu). We extracted perfectly conserved sequences from the whole genome alignments using a Python script available via a Git repository. To avoid false-positives (i.e. sequences aligned by pure chance), all perfectly conserved sequences under 5 base pairs were discarded. This threshold was computed based on an approach similar to the one in Eddy [2005]; the probability of two sequences end up perfectly aligned by chance rapidly decreases in function of their size (e.g. for one identical segment with 5 base pairs, the probability is around 0.00097 assuming a JC69 model).

### Windows

The human genome used as reference (hg38) was partitioned into non-overlapping consecutive windows of about 1,000 base pairs. During the definition of the window boundaries, we ensured that each PCS present in the 40 pairwise alignments would be entirely encompassed within a single window, thus avoiding the issue of cutting a PCS at the window’s edge. Due to this constraint, windows do not have exactly 1,000 base pairs, but approximately 1,000 base pairs.

### Estimates of Evolutionary Time

To assess the likelihood that the observed distribution of PCS sizes is associated with a specific evolutionary time *τ*, we begin by considering the number of PCSs present in the window. Then, for a given evolutionary time, we use Equation 2 to produce a sample of 100 PCS size distributions, ensuring that each sampled distribution matches the number of PCSs found in the observed data. We then use the Python library scipy.KDE to evaluate the likelihood that the observed data point is drawn from the same distribution that generated the 100 samples. Given its exponential-like shape, we calculated the logarithm of the PCS counts before comparing the PCS size distributions. This approach simplifies the estimations, as exponential functions appear linear when expressed logarithmically. The evolutionary time distribution of a window is then defined by aggregating the estimates derived from different evolutionary times. We applied this method for all windows in a parallelized way with the cluster computing resources provided by OIST.

### Identification of Functional Classes

All annotations for human (hg38) were obtained from UCSC Table Browser: coding sequence coordinates were taken from the tracks GENCODE-V44 and CCDS; repeats from the track RepeatMasker; and regulation elements from the RefSeq track. More information can be found in Supplementary Figure S7.

## Data and Software Availability

Analysis scripts and associated data reported in this article are archived as a Git repository through Zenodo at https://doi.org/XXXX (to be updated later).

## Acknowledgments

We thank Jules Lallouette and Jonathan Miller for providing a thorough review of this manuscript; Reuven Pnini, Lucia Zifcakova, Gabriela Cirtala and Simon Hellemans for helpful discussion; and OIST (Okinawa Institute of Science and Technology) Scientific Computing & Data Analysis Section for providing access to the high performance computing (HPC) and data storage resources. This study is supported by OIST core funding.

