## Supplementary Material for "Evolution of ultraconserved elements by indels"

Molecular Biology and Evolution

**Priscila Biller**

**Corresponding author: Priscila Biller.**
****

**This PDF file includes:**

Supporting text
Figs. S1 to S7
Tables S1 to S2
SI References

### Supporting Information Text

### Appendix A: Model of genome evolution by indels

In this section, we discuss the derivation of our model. Our model is an extension of the model proposed by Massip et al. (1, 2), which in turn is based on the pioneering work of Ziff and McGrady (3). Our model adds large mutations and mass loss to Massip's model, and it is based on a classical equation for discrete fragmentation:

$$\frac{\partial c_k(\tau)}{\partial \tau} = -a_k c_k(\tau) + \sum_{j=k+1}^{\infty} a_j b_{k|j} c_j(\tau). \qquad [1]$$

Applied to our case, the function $c_k(\tau)$ is defined as the expected number of perfectly conserved sequences (PCSs) of length $k$ at evolutionary time $\tau$. As discussed in the main text, evolutionary time $\tau$ is the product of mutation rate and chronological time, and should not be confused with the chronological time itself.

Consider a DNA sequence with $N$ base pairs. Initially in this process, at $\tau = 0$, both sequences are identical, thus $c_k(0)$ is 0 everywhere except at $k = N$, where $c_N(0)$ is equal to 1 (i.e. initially, both species/individuals share one sequence of size $N$). As the process continues, mutations accumulate, and this large PCS of size $N$ splits into smaller PCSs. The positions modified by mutations are not part of any PCS (in our case the process cannot be reversed), thus at $\tau > 0$, after mutations have occurred, we have:

$$0 \le \sum_{k=1}^{N} k c_k(\tau) < \sum_{k=1}^{N} k c_k(0) = N.$$

Here lies another difference between our model and Massip's model: whereas our model accounts for the mass loss caused by mutations, in Massip's model $\sum_{k=1}^{N} k c_k(\tau) = N$ during the whole process. This simplification is less of a problem in the "substitutions only" case, but it poses an issue for our model, where larger mutations can occur.

Next, we discuss the mutational process. Given one PCS of size $k$ shared by two individuals, mutations in either of the two copies can result in fragmentation of this PCS. A mutation of size $s$ that overlaps a PCS of size $k$ can take place in either $k + s - 1$ positions in one individual or $k + s - 1$ positions in the other, resulting in a total of $2(k + s - 1)$ possibilities. In our mutational process, mutations of any size can occur, and they may be fully contained within a PCS or partially overlap with it. The probability $P(s)$ of a mutation is inversely proportional to its size $s$ and is modeled as a truncated power-law (also called the Zipfian distribution). Putting these observations together, we can define the kernel $a_k$, which represents the breakup rate of a PCS of size $k$, as follows:

$$a_k = \sum_{s=1}^{\infty} 2(k+s-1)P(s) \propto \sum_{s=1}^{\infty} 2(k+s-1)s^{-\alpha} = 2((k-1)\zeta(\alpha) + \zeta(\alpha-1)),$$

where $\zeta(\alpha)$ is the Riemann zeta function and defined as $\zeta(\alpha) = \sum_{s=1}^{\infty} s^{-\alpha}$. As a side note, the computations are analogous if the mutation size is bounded to $N$. In this case, the terms $\zeta(\alpha)$ and $\zeta(\alpha - 1)$ in the preceding equation would be changed to $H_{N,\alpha}$ and $H_{N,\alpha-1}$, respectively. $H_{n,m}$ is the $n$th generalized harmonic number of order $m$ and defined as $H_{n,m} = \sum_{k=1}^{n} k^{-m}$.

We can use a similar idea to derive the other kernel $b_{k|j}$, which represents the average number of PCSs of size $k$ produced upon the breakup of a PCS of size $j$. As we saw earlier, if two individuals share a PCS of size $j$, then there are two identical sequences with $j$ base pairs, one for each individual. Looking at one of these two sequences, there are only two positions (left and right) where a mutation of size $s$ could occur to produce a PCS of size $k$. Thus, $a_j b_{k|j}$ can be written as:

$$a_j b_{k|j} = \sum_{s=1}^{\infty} 4P(s) \propto 4 \sum_{s=1}^{\infty} s^{-\alpha} = 4\zeta(\alpha),$$

where 4 is a result of the two possibilities (left and right) for each mutation size multiplied by two sequences, one for each individual.

To simplify our notation, let $\mu_{k,\alpha}$ be the mutation rate function, dependent on two parameters: the fragment size $k$ and the parameter $\alpha$, defined as

$$\mu_{k,\alpha} = 2((k-1)\zeta(\alpha) + \zeta(\alpha-1)).$$

By combining these results, we get

$$\frac{\partial c_k(\tau)}{\partial \tau} = -\mu_{k,\alpha} c_k(\tau) + \sum_{j=k+1}^{N} 4\zeta(\alpha) c_j(\tau). \qquad [2]$$

Equation 2 can be used to numerically compute evolutionary estimates. However, in our case, an analytical solution can also be derived using a similar approach to that of Huang et al. (4) (see their Appendix A).

Next, we proceed to solve this integro-differential equation using the Laplace transform method.

First, Equation 2 is differentiated with respect to $k$, and the summation is approximated by an integral to simplify the calculations:

$$\frac{\partial}{\partial k}\frac{\partial}{\partial \tau}c(k,\tau) = \frac{\partial}{\partial k}\Big(-\mu_{k,\alpha}c_k(\tau)\Big) + \frac{\partial}{\partial k}\left(\int_k^N 4\zeta(\alpha)c(j,\tau)\,dj\right), \qquad [3]$$

where $c_k(\tau) = c(k,\tau)$. We then fix $k$ and transform the $\tau$ variable using a Laplace transform. We denote by $s$ the transformed variable and $C(k,s)$ the transformed function:

$$C(k,s) = \mathcal{L}\{c(k,\tau)\} = \int_0^\infty c(k,\tau)e^{-s\tau}\,ds.$$

The transformation process for each term in Equation 3 will be outlined step by step, beginning with the leftmost term:

$$\mathcal{L}\left\{\frac{\partial}{\partial k}\frac{\partial}{\partial \tau}c(k,\tau)\right\} = \mathcal{L}\left\{\frac{\partial}{\partial \tau}\frac{\partial}{\partial k}c(k,\tau)\right\}$$

To transform the derivative in $\tau$ (the variable being transformed), we use the rule $\mathcal{L}\{y_t(x,t)\} = s\mathcal{L}\{y(x,t)\} - y(x,0)$. In our case, $y(x,t)$ is $\frac{\partial}{\partial k}c(k,\tau)$, and the transform of a derivative with respect to $k$ is equivalent to differentiating the transformed function:

$$\mathcal{L}\left\{\frac{\partial}{\partial k}c(k,\tau)\right\} = \int_0^\infty \frac{\partial}{\partial k}c(k,\tau)e^{-s\tau}\,ds = \frac{\partial}{\partial k}\left[\int_0^\infty c(k,\tau)e^{-s\tau}\,ds\right] = \frac{\partial}{\partial k}C(k,s).$$

Thus, the transformation of the leftmost term can be written as:

$$\mathcal{L}\left\{\frac{\partial}{\partial \tau}\frac{\partial}{\partial k}c(k,\tau)\right\} = s\mathcal{L}\left\{\frac{\partial}{\partial k}c(k,\tau)\right\} - \frac{\partial}{\partial k}c(k,0) = s\frac{\partial}{\partial k}C(k,s) - \frac{\partial}{\partial k}c(k,0), \qquad [4]$$

or $sC_k(k,s) - c_k(k,0)$ for a more concise representation.

Before proceeding with the Laplace transform of the next term in Equation 3, we will first make some simplifications:

$$\frac{\partial}{\partial k}\Big(-\mu_{k,\alpha}c(k,\tau)\Big) = -\left(2\zeta(\alpha)c(k,\tau) + \mu_{k,\alpha}\frac{\partial}{\partial k}c(k,\tau)\right),$$

which leads to

$$\mathcal{L}\left\{-\left(2\zeta(\alpha)c(k,\tau) + \mu_{k,\alpha}\frac{\partial}{\partial k}c(k,\tau)\right)\right\} = -2\zeta(\alpha)\mathcal{L}\{c(k,\tau)\} - \mu_{k,\alpha}\mathcal{L}\left\{\frac{\partial}{\partial k}c(k,\tau)\right\} \qquad [5]$$

$$= -2\zeta(\alpha)C(k,s) - \mu_{k,\alpha}\frac{\partial}{\partial k}C(k,s). \qquad [6]$$

The integral found in the rightmost term can be easily simplified thanks to the differentiation with respect to $k$:

$$\frac{\partial}{\partial k}\left(\int_k^N 4\zeta(\alpha)c(j,\tau)\,dj\right) = 4\zeta(\alpha)\frac{\partial}{\partial k}\Big(B(N,\tau) - B(k,\tau)\Big) = -4\zeta(\alpha)c(k,\tau).$$

Here, $B(k,\tau)$ refers to the antiderivative of $c(k,\tau)$—the usual notation $C(k,\tau)$ is not used to avoid confusion with the transformed function $C(k,s)$. Consequently, the transformation of the final term can be expressed simply as

$$\mathcal{L}\Big\{-4\zeta(\alpha)c(k,\tau)\Big\} = -4\zeta(\alpha)C(k,s). \qquad [7]$$

By combining the terms from Equations 4 to 7, we can define the transformation of Equation 3 as follows

$$s\frac{\partial}{\partial k}C(k,s) - \frac{\partial}{\partial k}c(k,0) = -2\zeta(\alpha)C(k,s) - \mu_{k,\alpha}\frac{\partial}{\partial k}C(k,s) - 4\zeta(\alpha)C(k,s). \qquad [8]$$

By rearranging the terms in Equation 8, we derive

$$(s+\mu_{k,\alpha})\frac{\partial}{\partial k}C(k,s) + 6\zeta(\alpha)C(k,s) = \frac{\partial}{\partial k}c(k,0) \qquad [9]$$

$$\frac{\partial}{\partial k}C(k,s) + \frac{6\zeta(\alpha)}{(s+\mu_{k,\alpha})}C(k,s) = \frac{1}{(s+\mu_{k,\alpha})}\frac{\partial}{\partial k}c(k,0). \qquad [10]$$

With the help of the Laplace transform, we obtain a first-order nonhomogeneous linear differential equation of the form $y' + p(k)y = f(k)$ (highlighted in gray). In our case

$$p(k) = \frac{6\zeta(\alpha)}{(s + \mu_{k,\alpha})},$$

and

$$f(k) = \frac{1}{(s + \mu_{k,\alpha})} \frac{\partial}{\partial k} c(k, 0).$$

As a result, Equation 10 can be solved using the integrating factor method. First, we calculate the integrating factor $I(k)$, defined as

$$I(k) = e^{\int p(k)dk}.$$

The integral of $p(k)$ is given by

$$\begin{aligned} \int p(k)dk &= \int \frac{6\zeta(\alpha)}{(s + 2((k-1)\zeta(\alpha) + \zeta(\alpha - 1)))} dk \\ &= 6\zeta(\alpha) \frac{1}{2\zeta(\alpha)} ln((s + 2((k-1)\zeta(\alpha) + \zeta(\alpha - 1)))) \\ &= 3ln(s + \mu_{k,\alpha}), \end{aligned}$$

and the integrating factor $I(k)$:

$$\begin{aligned} I(k) &= e^{\int p(k)dk} \\ &= e^{3ln(s+\mu_{k,\alpha})} \\ &= (e^{ln(s+\mu_{k,\alpha})})^3 \\ &= (s + \mu_{k,\alpha})^3 \end{aligned}$$

We continue by multiplying the entire equation by the integrating factor $I(k)$. The goal is to produce $I(k)y' + I'(k)y$ on the left side of Equation 10, which can be reformulated as $(I(k)y)'$ by applying the Leibniz product rule:

$$\begin{aligned} \frac{\partial}{\partial k} C(k, s) + \frac{6\zeta(\alpha)}{(s + \mu_{k,\alpha})} C(k, s) &= \frac{1}{(s + \mu_{k,\alpha})} \frac{\partial}{\partial k} c(k, 0) \\ (s + \mu_{k,\alpha})^3 \frac{\partial}{\partial k} C(k, s) + \frac{6\zeta(\alpha)(s + \mu_{k,\alpha})^3}{(s + \mu_{k,\alpha})} C(k, s) &= \frac{(s + \mu_{k,\alpha})^3}{(s + \mu_{k,\alpha})} \frac{\partial}{\partial k} c(k, 0) \\ (s + \mu_{k,\alpha})^3 \frac{\partial}{\partial k} C(k, s) + 6\zeta(\alpha)(s + \mu_{k,\alpha})^2 C(k, s) &= (s + \mu_{k,\alpha})^2 \frac{\partial}{\partial k} c(k, 0) \\ \frac{\partial}{\partial k} \Big( (s + \mu_{k,\alpha})^3 C(k, s) \Big) &= (s + \mu_{k,\alpha})^2 \frac{\partial}{\partial k} c(k, 0). \end{aligned}$$

By integrating both sides of the equation, a solution for $C(k, s)$ can be obtained as follows:

$$\int^k \frac{\partial}{\partial u} \Big( (s + \mu_{u,\alpha})^3 C(u, s) \Big) \partial u = \int^k \Big( (s + \mu_{u,\alpha})^2 \frac{\partial}{\partial u} c(u, 0) \Big) \partial u,$$

the left side of the equation can be simplified easily, while the right side can be solved using the integration by parts rule, defined as $\int f(k)g'(k)dk = f(k)g(k) - \int g(k)f'(k)dk$. In our case, we assign

$$\begin{aligned} f(u) &= (s + \mu_{u,\alpha})^2 \\ f'(u) &= 4\zeta(\alpha)(s + \mu_{u,\alpha}) \\ g(u) &= c(u, 0) \\ g'(u) &= \frac{\partial}{\partial u} c(u, 0), \end{aligned}$$

which leads to:

$$(s+\mu_{k,\alpha})^3 C(k,s) = (s+\mu_{k,\alpha})^2 c(k,0) - \int^k \Big( c(u,0) 4\zeta(\alpha)(s+\mu_{u,\alpha}) \Big) \partial u.$$

By separating the term $C(k,s)$ to the left side, we obtain a solution for the transformed function $C(k,s)$:

$$C(k,s) = \frac{c(k,0)}{(s+\mu_{k,\alpha})} - \frac{4\zeta(\alpha)}{(s+\mu_{k,\alpha})^3} \int^k \Big( c(u,0)(s+\mu_{u,\alpha}) \Big) \partial u. \tag{11}$$

We are now set to apply the inverse Laplace transform to $C(k,s)$ to find a solution for the original function $c(k,\tau)$. We will describe the transformation process for every term in Equation 11 step by step, beginning with the leftmost term, which is straightforwardly:

$$\mathcal{L}_\tau^{-1}\Big\{ C(k,s) \Big\} = c(k,\tau). \tag{12}$$

Continuing with the next term in Equation 11, we are going to use the rule $\mathcal{L}_\tau^{-1}\Big\{ \frac{1}{s+x} \Big\} = e^{-\tau x}$, by setting $x = \mu_{k,\alpha}$:

$$\mathcal{L}_\tau^{-1}\Big\{ \frac{c(k,0)}{(s+\mu_{k,\alpha})} \Big\} = c(k,0) e^{-\tau \mu_{k,\alpha}}. \tag{13}$$

Before we apply the inverse Laplace transform to the rightmost term of Equation 11, we will first take out the variable $s$ from the integral:

$$-\frac{4\zeta(\alpha)}{(s+\mu_{k,\alpha})^3} \int^k \Big( c(u,0)(s+\mu_{u,\alpha}) \Big) \partial u = -\frac{4\zeta(\alpha)s}{(s+\mu_{k,\alpha})^3} \int^k c(u,0) \partial u \tag{14}$$

$$- \frac{4\zeta(\alpha)}{(s+\mu_{k,\alpha})^3} \int^k \Big( c(u,0)\mu_{u,\alpha} \Big) \partial u. \tag{15}$$

$$\tag{16}$$

Using the fact that

$$\mathcal{L}_\tau^{-1}\Big\{ \frac{s}{(s+a)^3} \Big\} = e^{-a\tau} \tau \Big( 1 - \frac{a\tau}{2} \Big)$$

$$\mathcal{L}_\tau^{-1}\Big\{ \frac{1}{(s+a)^3} \Big\} = e^{-a\tau} \tau^2 \frac{1}{2},$$

we obtain:

$$\mathcal{L}_\tau^{-1}\Big\{ -\frac{4\zeta(\alpha)s}{(s+\mu_{k,\alpha})^3} \int^k c(u,0)\partial u \Big\} = -4\zeta(\alpha) \int^k c(u,0)\partial u \; \mathcal{L}_\tau^{-1}\Big\{ \frac{s}{(s+\mu_{k,\alpha})^3} \Big\} \tag{17}$$

$$= -4\zeta(\alpha) \int^k c(u,0)\partial u \; e^{-\tau\mu_{k,\alpha}} \; \tau \Big( 1 - \frac{\tau\mu_{k,\alpha}}{2} \Big) \tag{18}$$

$$\mathcal{L}_\tau^{-1}\Big\{ -\frac{4\zeta(\alpha)}{(s+\mu_{k,\alpha})^3} \int^k \Big( c(u,0)\mu_{u,\alpha} \Big) \partial u \Big\} = -4\zeta(\alpha) \int^k c(u,0)\mu_{u,\alpha}\partial u \; \mathcal{L}_\tau^{-1}\Big\{ \frac{1}{(s+\mu_{k,\alpha})^3} \Big\} \tag{19}$$

$$= -4\zeta(\alpha) \int^k c(u,0)\mu_{u,\alpha}\partial u \; e^{-\tau\mu_{k,\alpha}} \tau^2 \frac{1}{2} \tag{20}$$

Combining all calculated terms (equations 12–20), we can derive the inverse Laplace transform of Equation 11:

$$\begin{aligned} c(k,\tau) &= c(k,0)e^{-\tau\mu_{k,\alpha}} \\ &- 4\zeta(\alpha) \int^k c(u,0)\partial u \; e^{-\tau\mu_{k,\alpha}} \; \tau \Big( 1 - \frac{\tau\mu_{k,\alpha}}{2} \Big) \\ &- 4\zeta(\alpha) \int^k c(u,0)\mu_{u,\alpha}\partial u \; e^{-\tau\mu_{k,\alpha}} \tau^2 \frac{1}{2}, \end{aligned}$$

which can be simplified further to

$$\begin{aligned} c(k,\tau) &= e^{-\tau\mu_{k,\alpha}} \Big[ c(k,0) + 4\zeta(\alpha) \Big( \tau \Big( 1 - \frac{\tau\mu_{k,\alpha}}{2} \Big) \int_k^\infty c(u,0)\partial u + \frac{\tau^2}{2} \int_k^\infty c(u,0)\mu_{u,\alpha}\partial u \Big) \Big] \\ &= e^{-\tau\mu_{k,\alpha}} \Big[ c(k,0) + 4\zeta(\alpha) \int_k^\infty c(u,0) \frac{\tau}{2} \Big( 2 + \tau(\mu_{u,\alpha} - \mu_{k,\alpha}) \Big) \partial u \Big], \end{aligned}$$

thus providing the following solution for $c(k,\tau)$:

$$c(k,\tau) = e^{-\tau\mu_{k,\alpha}}\left[c(k,0) + 2\tau\zeta(\alpha)\int_k^\infty c(u,0)[2+\tau(\mu_{u,\alpha}-\mu_{k,\alpha})]\partial u\right]. \quad [21]$$

Equation 21 is similar to Equation 11 given by Ziff and Grady (3). This equation can be further simplified by incorporating the initial conditions of the evolutionary process. At time $\tau = 0$, all sequences are identical, and individuals share one PCS of size $N$:

$$c(k,0) = \begin{cases} 1, & \text{if } k = N \\ 0, & \text{if } 0 < k < N. \end{cases} \quad [22]$$

Therefore, the integral term in Equation 21 is simplified to

$$c(k,\tau) = \begin{cases} e^{-\tau\mu_{N,\alpha}}(1+4\tau\zeta(\alpha)), & \text{if } k = N \\ e^{-\tau\mu_{k,\alpha}}(4\tau\zeta(\alpha)+2\tau^2\zeta(\alpha)(\mu_{N,\alpha}-\mu_{k,\alpha})), & \text{if } 0 < k < N. \end{cases}$$

which was the equation used to compute the evolutionary estimates discussed in the text.

The model proposed by Massip and Arndt (1, 2) encompasses substitutions only, which is equivalent to setting the parameter $\alpha$ to a very high value in our model. When $\alpha \to \infty$, the Riemann zeta function $\zeta(\alpha)$ approaches 1 and the mutation function $\mu_{k,\alpha}$ is equal to $2((k-1)\zeta(\alpha)+\zeta(\alpha-1)) = 2(k-1)1+1 = 2k$. Thus, for the general case $0 < k < N$, we have:

$$\begin{aligned} c(k,\tau) &= e^{-\tau\mu_{k,\alpha}}(4\tau\zeta(\alpha)+2\tau^2\zeta(\alpha)(\mu_{N,\alpha}-\mu_{k,\alpha})) \\ &= e^{-\tau 2k}(4\tau+2\tau^2(2N-2k)) \\ &= e^{-\tau 2k}(4\tau+4\tau^2(N-k)), \end{aligned} \quad [23]$$

which is exactly Equation 5 (setting $\mu = 1$) in Massip and Arndt's work (2).

### Appendix B: Dataset details

**Table S1. List of genomes used in our analysis (assembly IDs and divergence times), sorted by divergence time. Divergence time is displayed in million of years ago (mya). All pairwise alignments between human (hg38) and another vertebrate were downloaded from the UCSC website (http://genome.ucsc.edu/, last accessed in Feb./2025). 40 pairwise alignments were used in our analysis (i.e. all available pairwise alignments involving the human genome assembly "hg38" at the time). Divergence times were taken from TimeTree (http://timetree.org/, last accessed in Feb./2025), except for close primates (bonobo, chimp, and gorilla, marked with $\star$), whose information was taken from a more recent study (5) (see also (6)).**

| Species | Assembly ID | Divergence time (mya) |
|---|---|---:|
| 01. Bonobo | panPan3 | $\star$12.1 |
| 02. Chimp | panTro6 | $\star$12.1 |
| 03. Gorilla | gorGor6 | $\star$15.1 |
| 04. Orangutan | ponAbe3 | 15.2 |
| 05. Baboon | papAnu4 | 28.8 |
| 06. Crab-eating macaque | macFas5 | 28.8 |
| 07. Golden snub-nosed monkey | rhiRox1 | 28.8 |
| 08. Green monkey | chlSab2 | 28.8 |
| 09. Proboscis Monkey | nasLar1 | 28.8 |
| 10. Rhesus | rheMac10 | 28.8 |
| 11. Marmoset | calJac4 | 43.0 |
| 12. Tarsier | tarSyr2 | 69.0 |
| 13. Mouse lemur | micMur2 | 74.0 |
| 14. Malayan flying lemur | galVar1 | 79.0 |
| 15. Mouse | mm39 | 87.0 |
| 16. Rabbit | oryCun2 | 87.0 |
| 17. Rat | rn7 | 87.0 |
| 18. Alpaca | vicPac2 | 94.0 |
| 19. Bison | bisBis1 | 94.0 |
| 20. Cat | felCat9 | 94.0 |
| 21. Chinese pangolin | manPen1 | 94.0 |
| 22. Cow | bosTau9 | 94.0 |
| 23. Dog | canFam6 | 94.0 |
| 24. Ferret | musFur1 | 94.0 |
| 25. Hawaiian monk seal | neoSch1 | 94.0 |
| 26. Horse | equCab3 | 94.0 |
| 27. Little brown bat | myoLuc2 | 94.0 |
| 28. Pig | susScr11 | 94.0 |
| 29. Southern sea otter | enhLutNer1 | 94.0 |
| 30. Manatee | triMan1 | 99.0 |
| 31. Wallaby | macEug2 | 160.0 |
| 32. Platypus | ornAna2 | 180.0 |
| 33. Brown kiwi | aptMan1 | 319.0 |
| 34. Chicken | galGal6 | 319.0 |
| 35. Garter snake | thaSir1 | 319.0 |
| 36. Golden eagle | aquChr2 | 319.0 |
| 37. Turkey | melGal5 | 319.0 |
| 38. African clawed frog | xenLae2 | 352.0 |
| 39. X. tropicalis | xenTro10 | 352.0 |
| 40. Zebrafish | danRer11 | 429.0 |

### Appendix C: Evolutionary time estimates—Effect of treating fast-evolving and slow-evolving regions as a single region

Our method relies on the assumption of uniform evolution across the window. In this section we discuss in more details the case where the analyzed region is composed of subregions with very distinct evolutionary times, i.e., when fast-evolving and slow-evolving regions are mixed together and treated as one single region.

The first important observation is that, during the evolutionary process, the number of PCSs increases as mutations accumulate in the sequence. The number of mutations is related to the evolutionary time $t$: the higher the number of mutations is, the higher the value of $t$ is (Figure S5, panel A). The evolutionary time $t$ is the product of chronological time and mutation rate. Two adjacent regions evolving at different evolutionary times must have different mutation rates, as the chronological time must be identical. If two adjacent regions are evolving with different mutation rates, and their PCS size distributions are mixed together, then the combined PCS size distribution will contain a higher number of PCSs coming from the fast-evolving region (Figure S5, panel B). For example, in an extreme case, one single long PCS coming from a very conserved subregion could end up mixed with $n$ small PCSs coming from one fast-evolving subregion.

The method used to estimate $t$ takes as input this mixed PCS size distribution, and finds the evolutionary time whose expected PCS size distribution is the closest to the observed (mixed) PCS size distribution. The "closest distribution" can be computed in different ways. One possible way is to define the closest distribution as the one that minimizes the difference between observed and estimated counts along the distribution. Since the mixed PCS size distribution is mainly composed of PCSs from the fast evolving region, the counts are dominated by the fast evolving region, biasing the method that selects $t$, as shown in Figure S5 (panel C). To reduce this bias, we chose a relatively small window size (1,000 bps), which hopefully minimizes the heterogeneity of the mutation rate within the window. The window size, however, cannot be too small; otherwise the window might not have enough information for the estimation.

### Appendix D: Mutation rate in fast-evolving regions

For Fig. 3 in the main text, panel B (left graph), the mutation rate $\mu$ was set at $18 \times 10^{-9}$ per position per year (PPPY). The estimated mutation rate for fast-evolving regions is based on the values found in the high tail ($>$ 99th percentile) of estimated evolutionary time distributions. As a reference point, this value is about eight times higher than the average mutation rate found in mammals in previous research (10). This value is expected to be significantly higher than the average, as it reflects the mutation rate solely in regions where the PCS size distributions are no longer informative.

**Table S2. Indel rates from previous works. Method: "Direct estimates" (D) refers to methods that count mutations that occur between generations in present-day individuals, whereas "Indirect estimates" (I) refers to estimates based on the evolutionary distance separating two species divided by (twice) their divergence time. Divergence time was taken from different sources, indicated in each row at column "Observations". Mutation info: size range of mutations considered in the analysis (in base pairs), and their type (insertions, deletions, or both). Mutation rate estimates: original value reported in previous works, and the adapted value (converted to PPPY) used in our analysis. Mutation rate units: PPPG = Per Position Per Generation; PPPY = Per Position Per Year; PPPL = Per Position Per Lineage. (*) In Lunter's work, the referred lineage is the one connecting human and mouse. (**) In Nachman and Crowell's work, the referred lineage is the one connecting human and chimpanzee. The value 0.0012 was taken from Table 1 ("Rates of evolution for autosomal processed pseudogenes"), column $K_i$, row "Mean autosomal values".**

| Study | Method | Mutation info | | | Mutation rate estimates | | Observations |
|---|---|---|---|---|---|---|---|
| | | Size (bp) | Ins. | Del. | Original value | Adapted (PPPY) | |
| 1. Maretty et al. (2017) (11) | D | $\leq 10$ | $\star$ | $\star$ | $1.3 \times 10^{-9}$ PPPG | $4.7 \times 10^{-11}$ | 1 generation = 27.7 years (provided in their study) |
| 2. Besenbacher et al. (2016) (12) | D | $\leq 35$ | $\star$ | $\star$ | $0.929^{-9} \times 10^{-9}$ PPPG | $3.07 \times 10^{-11}$ | Their study provides indel rates both in PPPG and PPPY (est. generation time: 30.26 years). |
| 3. Kloosterman et al. (2015) (13) | D | $\leq 20$ | $\star$ | $\star$ | $0.68 \times 10^{-9}$ PPPG | $2.32 \times 10^{-11}$ | 1 generation = 29.27 years (based on the average age of 87 individuals reported in their supplementary material) |
| 4. Palamara et al. (2015) (14) | D | $\leq 20$ | $\star$ | $\star$ | $1.26 \times 10^{-9}$ PPPG | $4.34 \times 10^{-11}$ | 1 generation = 29 years (value assumed in their paper for converting PPPG to PPPY) |
| 5. Besenbacher et al. (2015) (15) | D | $\leq 50$ | $\star$ | $\star$ | $1.5 \times 10^{-9}$ PPPG | $5.28 \times 10^{-11}$ | 1 generation = 28.4 years (provided in their study, only fathers taken into account) |
| 6. Kondrashov (2002) (16) | D | $\leq 5$ | $\star$ | | $0.182 \times 10^{-9}$ PPPG | $0.91 \times 10^{-11}$ | 1 generation = 20 years (suggested value in his paper). Maximum mutation size is based on Figure 5 from Kondrashov's paper. |
| 7. Kondrashov (2002) (16) | D | $\leq 5$ | | $\star$ | $0.526 \times 10^{-9}$ PPPG | $2.63 \times 10^{-11}$ | 1 generation = 20 years (suggested value in his paper). Maximum mutation size is based on Figure 5 from Kondrashov's paper. |
| 8. Lunter (2007) (17) | I | any | $\star$ | $\star$ | (*) 0.053 PPPL | $30.46 \times 10^{-11}$ | Time separating human-mouse = $2 \times 87 \times 10^6$ years. Source of divergence time: TimeTree (87 mya). |
| 9. Nachman and Crowell (2000) (18) | I | $\leq 4$ | $\star$ | $\star$ | (**) 0.0012 PPPL | $4.95 \times 10^{-11}$ | Time separating human-chimpanzee = $2 \times 12.1 \times 10^6$ years. Source of divergence time: Moorjani et al. (2016) (12.1 mya)(5). |

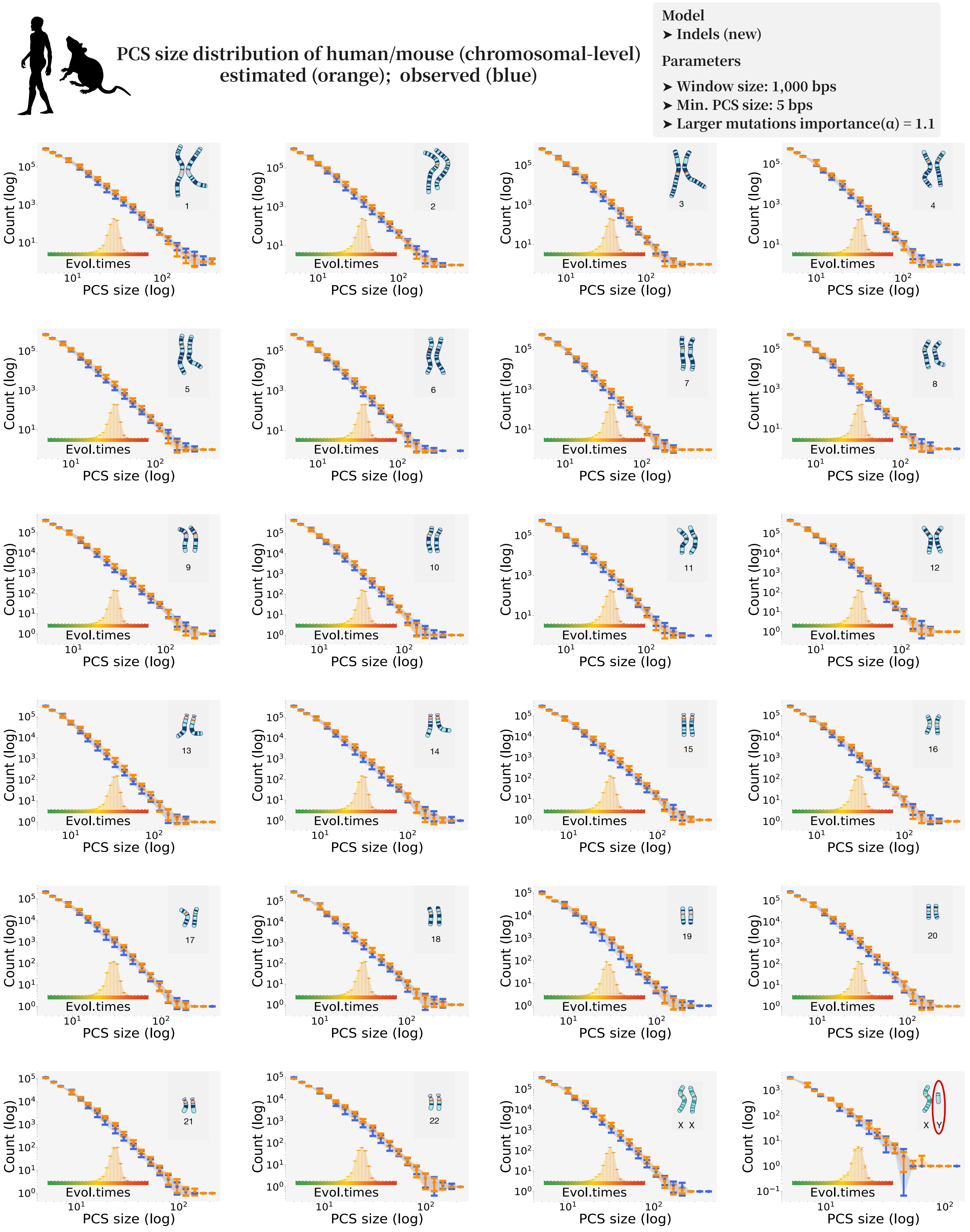


**Fig. S1. Evolutionary estimates from our model: Indels / Chromosome level.** Length distribution of ultraconserved elements shared by human and mouse. Perfectly conserved sequence (PCS) size distributions are grouped by human chromosome. The corresponding chromosome is indicated at the top right of each plot. Observed data are shown in blue, and predictions from our model are shown in orange.

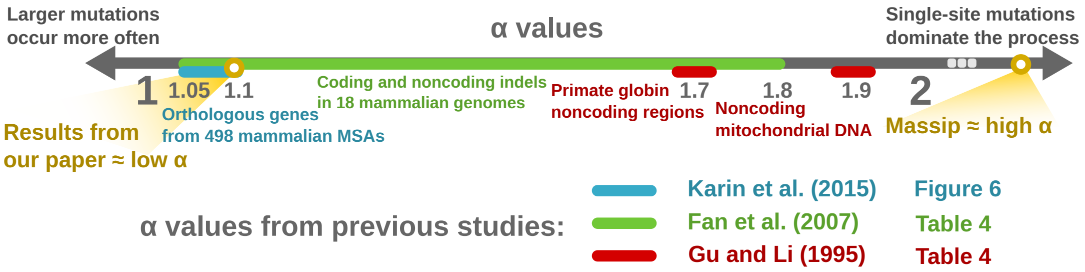


**Fig. S2.** $\alpha$ **values inferred in previous works.** Zipfian distributions have been largely used to model the probability of insertions and deletions based on their length. The value of its single parameter $\alpha$ has been previously inferred for different genomic datasets. Some examples are shown in this figure: blue: Karin et al. (2015) (7); green: Fan et al. (8); red: Gu and Li (1995) (9).

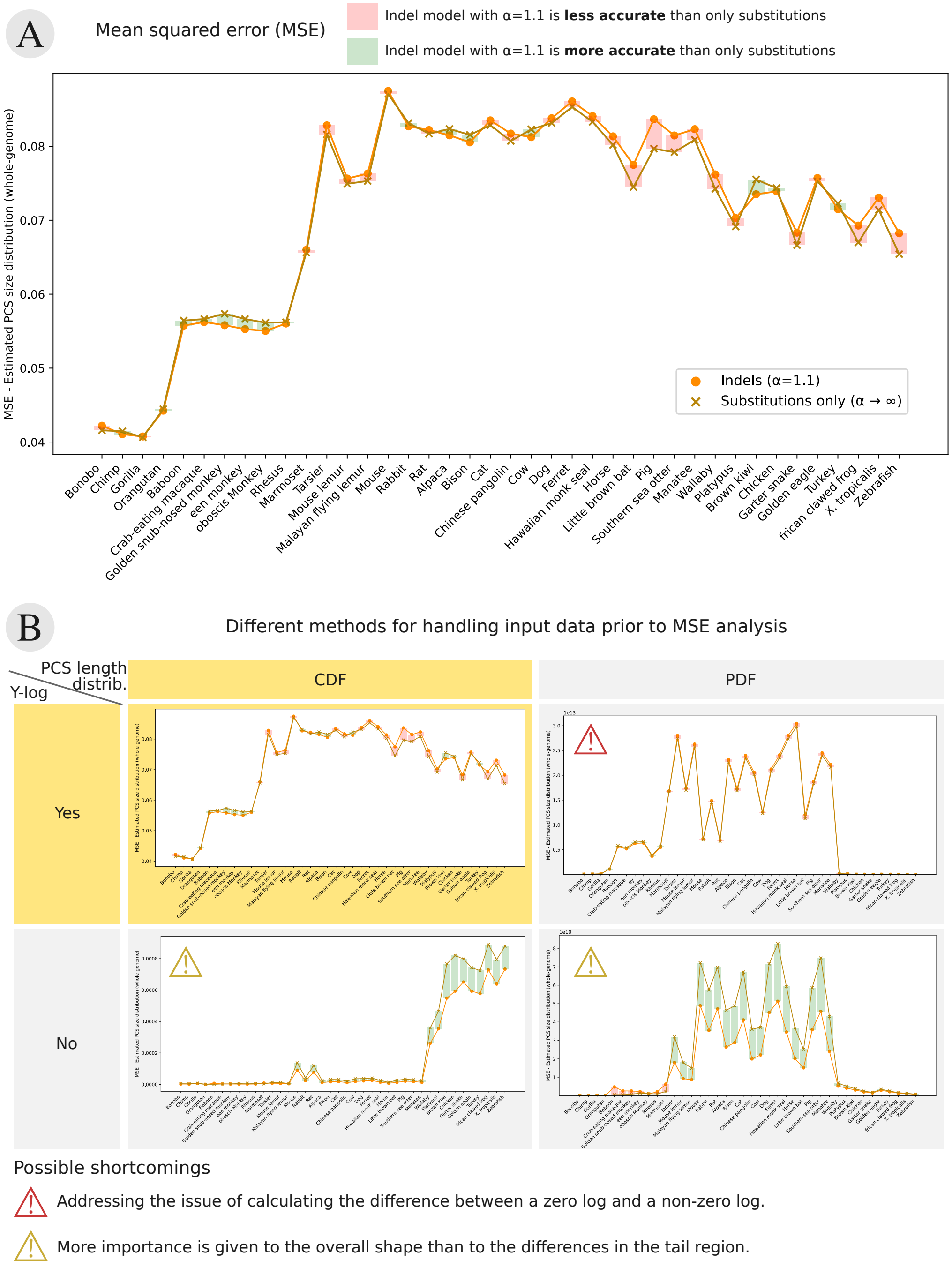


**Fig. S3. Mean squared error (MSE).** Comparison of the MSE for two evolutionary models: a model with large events (light orange markers, parameter $\alpha$ is set to $1.01$ for all cases) and a model with substitutions only (dark yellow markers, Massip et al. (2015) (1)). The graph shows the error (y-axis) between estimated and observed PCS size distributions at whole-genome level for each species (x-axis; species are sorted by divergence time, from the species closest to human on the left to the farthest species on the right). The green (red) areas highlight the cases where a model with large events is more (less) accurate than a model with substitutions only. The difference between the methods lies in the estimated counts of large PCSs (i.e. the tail of the PCS length distribution). To effectively account for the discrepancies in the tail of the distribution, two important considerations should be noted: (1) Large PCSs are found dispersed throughout a wide range of sizes, and (2) their counts are orders of magnitude lower than those of small PCSs. The former was handled by binning the observed and estimated PCS size distributions prior to calculating the MSE, whereas the latter was tackled by taking the logarithm of the counts for each bin. **Upper graph, marked as (A)**, shows the resulting MSE from the cumulative distribution (CDF) of the logarithmic counts. The choice of CDF is linked to the use of a logarithmic scale: as bins encompassing large PCS sizes might be empty, the difference of the logarithmic counts can become ill-defined, and CDF helps to avoid this scenario. **Lower graphs, marked as (B)**, show different methods of pre-processing the distributions before MSE computation, along with the resulting MSE. The top and bottom rows display the resulting MSE in logarithmic and linear scales, while the left and right columns present the MSE from CDF and PDF, respectively. The graph highlighted in yellow in (B) corresponds to the graph in (A).

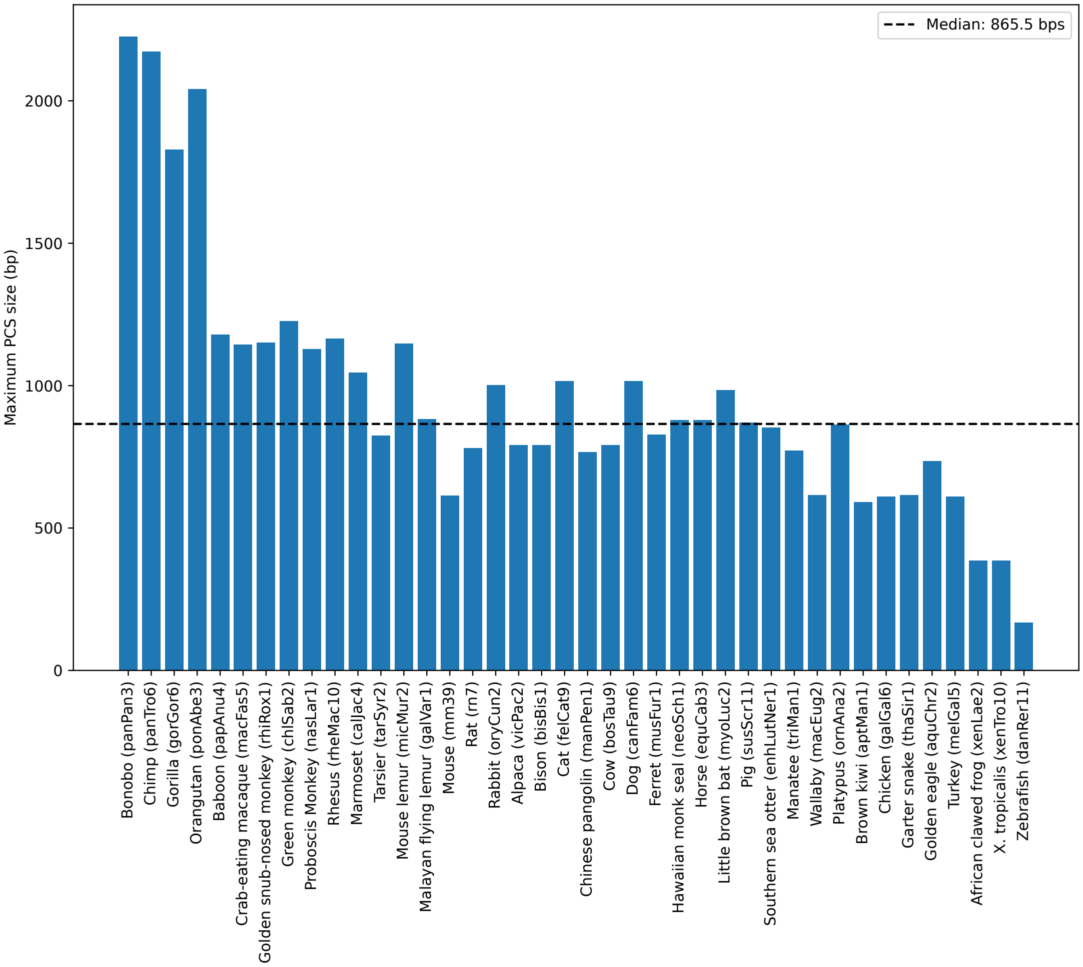


**Fig. S4. Maximum size of perfectly conserved sequences (PCS).** Size of the largest perfectly conserved sequence (y-axis, shown in base pairs) found in the pairwise alignment between human and another species (x-axis, sorted by divergence time from human).

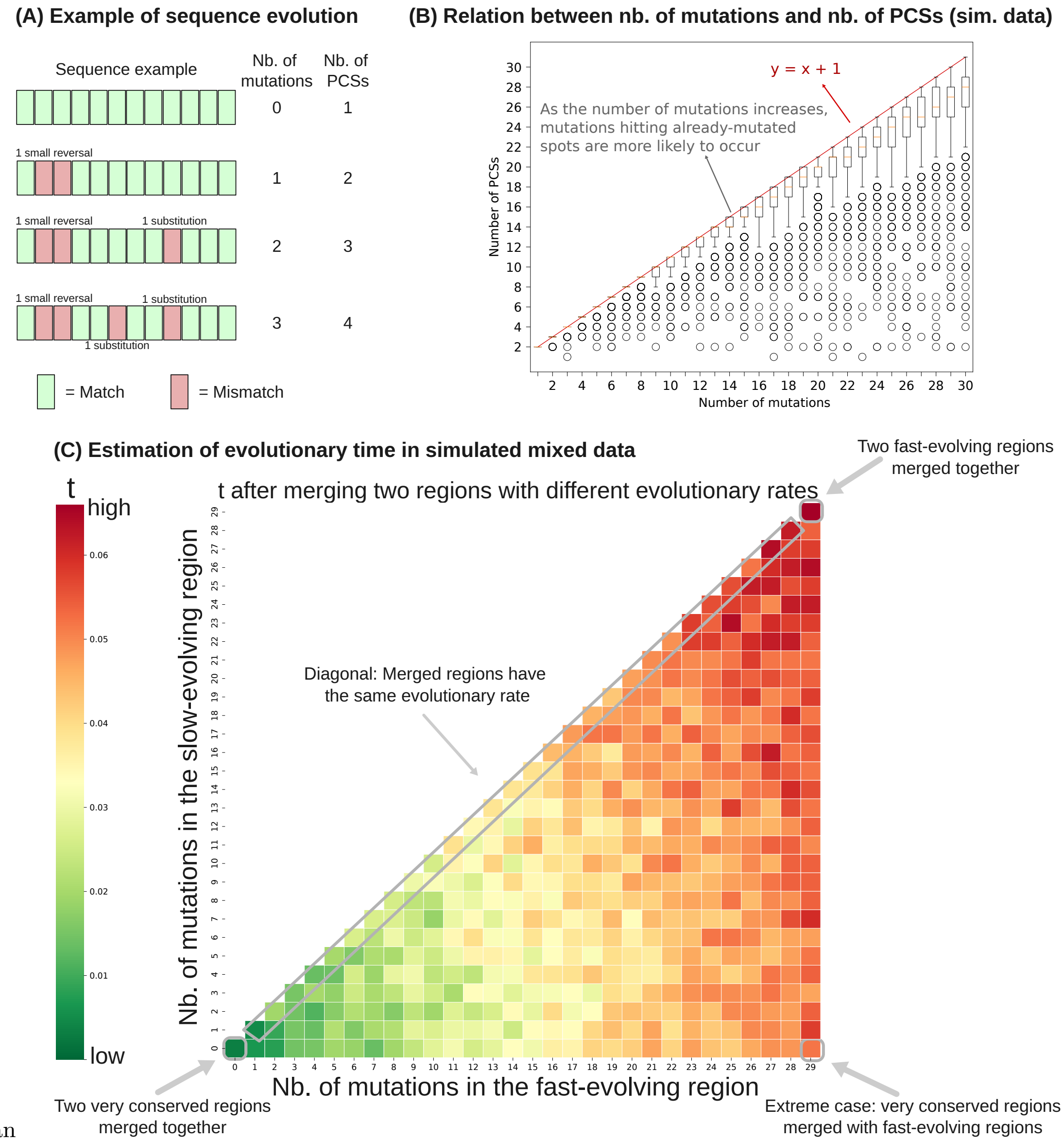


than

**Fig. S5. Effect of mixing fast-evolving and slow-evolving regions together. Panel A:** Schematics showing how the number of PCSs changes as mutations accumulate in the sequence. Green rectangles are matches between two sequences, whereas red rectangles are mismatches or gaps (i.e. a mutation occurred in either one of the sequences). **Simulated data in panels B and C:** a sequence with 1,000 positions was evolved by a simple evolutionary process consisting of applying one mutation at every step. Mutations are randomly chosen by uniformly sampling one position between $1 \ldots 1000$, and one mutation size from a Zipfian distribution ($\alpha$=2). This process was repeated 1,000 times. **Panel B:** Relation between the number of PCSs (y-axis) and the number of mutations (x-axis) on simulated datasets. **Panel C:** Estimated evolutionary time when fast-evolving and slow-evolving regions are combined. The method overestimates evolutionary times in scenarios where the uniform assumption is invalid.

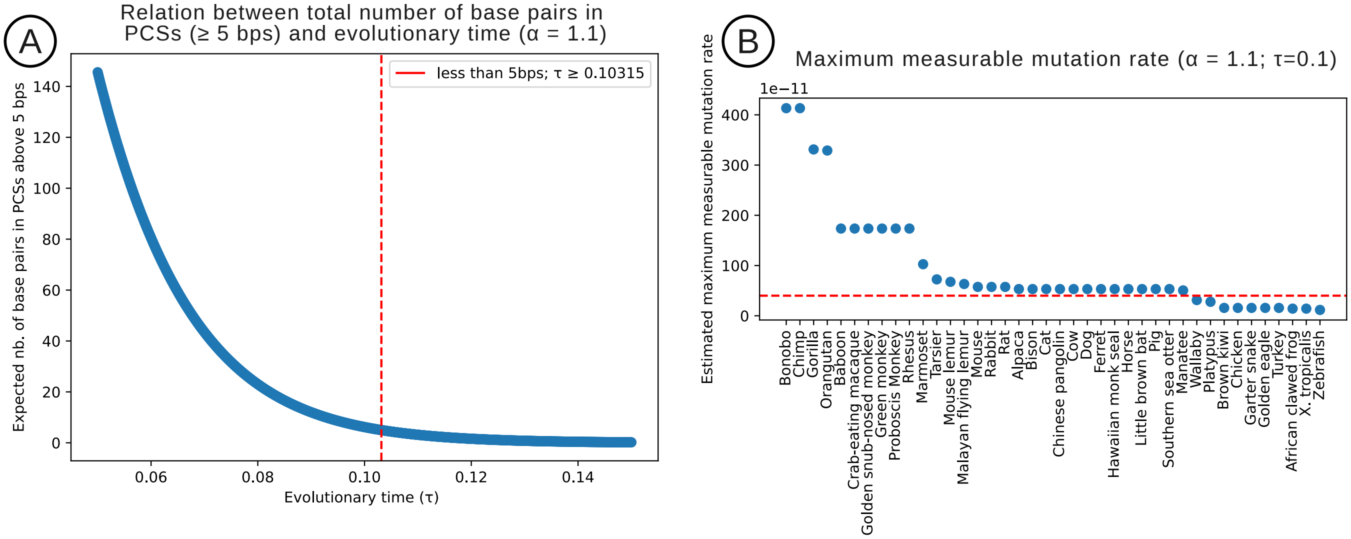


**Fig. S6.** Maximum measurable mutation rate. **Panel A (left):** Expected number of base pairs in PCSs (y-axis) depending on the evolutionary time $\tau$ (x-axis), in a model where $\alpha$ was set to $1.1$ (indels). The values shown in the y-axis were computed with Equation **??**. **Panel B (right):** Maximum indel rate (y-axis, values given in per position per year) that can be inferred with our model based on PCSs, for each species in our dataset (x-axis). Given that mutation rates in rapidly evolving regions can exceed $50 \times 10^{-11}$ PPPY, we expect that species with maximum mutation rates below the red dashed line are likely to have their indel rates underestimated.

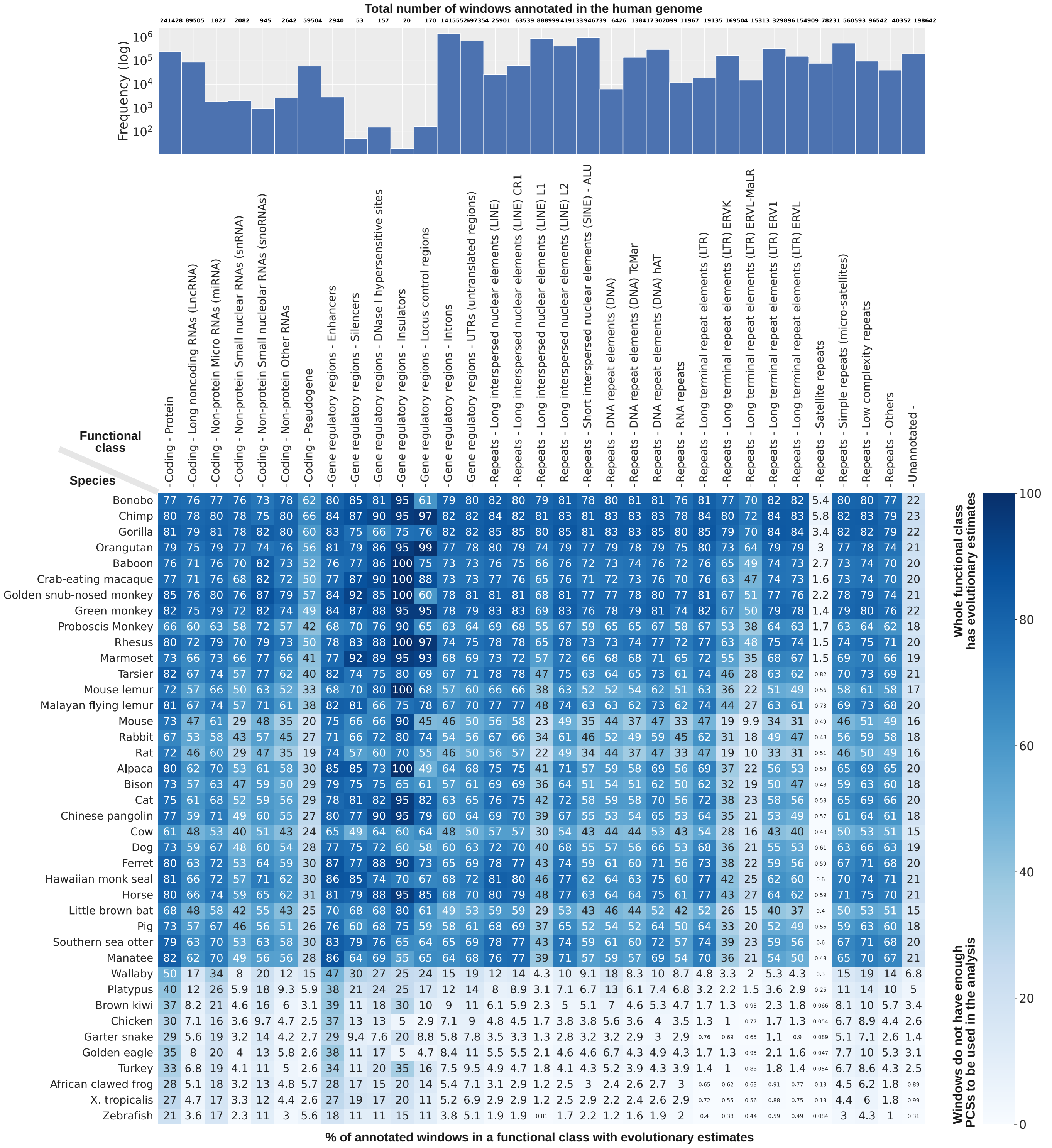


**Fig. S7. Functional classes. Top graph:** Total number of windows (y-axis, log scale) annotated per functional class (x-axis), taking into account both windows used in our analysis and windows ignored in our analysis. Windows used in our analysis are the ones with "enough" information to have an evolutionary estimate, i.e., the window must have at least one PCS above 8 base pairs. Windows ignored in our analysis are the ones fast-evolving, containing only PCSs below 8 base pairs, making it not possible to use the curve-fitting method described in the main text. **Bottom graph:** Percentage of the total number of windows per functional class (y-axis) that have an evolutionary estimate. Rows show the species sorted by divergence time. As expected, as divergence time increases, less windows have enough PCSs to be used in the analysis. Percentages below 1% are displayed in a smaller font to be able to fit in the cell.
